# A robust platform for recombinant production of animal venom toxin modulators of ion channels

**DOI:** 10.1101/2025.06.11.659050

**Authors:** Jose Enrique Gonzalez-Prada, Alexander S. Haworth, Jacob I. Browne, Samantha C. Salvage, Nicola Secomandi, Anthony M. Rush, Anne Ritoux, Maya Dannawi, Leonard Lee, Vasiliki Mavridou, Yin Y. Dong, Ewan SJ. Smith, Antony P. Jackson, Edward B. Stevens, Paul S. Miller

**Affiliations:** Department of Pharmacology, University of Cambridge, Cambridge, UK; Metrion Biosciences, Granta Park, Cambridge, UK; Department of Biochemistry, University of Cambridge, Cambridge, UK; St George’s School of health and medical sciences, City St George’s, London, UK; Nuffield Department of Clinical Neurosciences, University of Oxford, Oxford, UK

**Keywords:** Automated patch-clamp, electrophysiology, immunocytochemistry, peptides, toxin, protein expression, recombinant, pain target, protoxin II, huwentoxin IV, pull-down, voltage-gated sodium channel, nAChR, ion channel

## Abstract

**Background and Purpose:** Peptide toxins isolated from animal venom are potent and selective modulators of ion channels, and promising therapeutic leads. Due to intricate disulphide bridge patterns, they are often challenging to produce in standard laboratory settings, which limits engineering approaches to manipulate their structure-function properties. Given the low cost, wide accessibility, and versatility of recombinant expression systems for protein production, we set out to establish a straightforward high-yield strategy across a broad panel of peptide toxins from snakes, spiders and scorpions.

**Experimental Approach:** 13 toxin DNA sequences were genetically fused to the C-terminus of either bivalent or monovalent human IgG1 antibody fragment crystallisable (Fc) domain sequences and expressed recombinantly from mammalian Expi293F cells. Affinity-purified proteins were evaluated by SDS-PAGE and size-exclusion chromatography (SEC). Function was assessed by Ca^2+^ flux assays on CN21 cells, or whole-cell electrophysiology on human embryonic kidney (HEK293T) cells, Chinese hamster ovary (CHO) cells, or dorsal root ganglion (DRG) neurons. Immunocytochemistry using HEK293T cells and mouse DRG neurons assessed Fc-toxin fusion binding.

**Key Results:** Monovalent Fc-toxin fusions consistently yielded 1-6 mg of pure, non-proteolytically cleaved protein from 20-70 ml cultures for several toxin types, including three-finger toxins from snakes, inhibitory cystine knot (ICK) toxins from spiders, and α-toxins from scorpions, substantially surpassing the performance of unfused toxins or bivalent Fc-toxin fusions which gave low or no yield. Snake toxins targeting nicotinic acetylcholine receptors retained high single digit nanomolar inhibitory potency. Spider and scorpion toxins targeting the voltage-gated Na^+^ channel Na_v_1.7 retained pharmacological function and selectivity across a panel of five Na_v_ subtypes, albeit with reduced potencies that did not exceed ∼70 nM.

**Conclusions and Implications:** We present a strategy for straightforward robust production of pure, monodisperse, and functional animal venom-derived toxins. This lowers the barrier to toxin production in a standard laboratory setting for follow-on engineering purposes.

## INTRODUCTION

Animal venom has proven to be a rich source of novel peptide toxins with varied pharmacological properties (Herzig et al., 2020; Cardoso et al., 2022). Many are potent and selective modulators of ion channels, such as nicotinic acetylcholine receptors (nAChRs) and voltage-gated Na^+^ channels (Na_v_s) (Kalia et al., 2015; Cardoso and Lewis, 2018; Wulff et al., 2019; Robinson and Vetter, 2020). Given that ion channels are high-value drug targets and achieving small molecule selectivity remains challenging, peptide toxins have sparked considerable interest to advance both basic research and scaffold-based drug discovery (Montnach et al., 2021; Nguyen et al., 2022).

Many of the best characterised venom-derived peptide toxins possess intricate arrangements of disulphide bridges. These include, for example, snake venom three-finger toxins, such as α-cobratoxin (αCbTx) (Bourne et al., 2005), spider venom toxins that harbour inhibitory cystine knot (ICK) motifs, including protoxin-II (ProTxII) (Xu et al., 2019), and scorpion venom α-toxins, like AahII (Clairfeuille et al., 2019). The disulphide bridges endow peptide toxins with high biochemical stability but constitute a barrier to production from recombinant cell expression platforms due to disulphide scrambling causing misfolding and loss of potency relative to synthetically produced toxins. Typical recombinant production approaches involve fusing toxins to carrier proteins that enhance solubility, hence improving yields, and/or prolonging toxin plasma half-life for *in vivo* applications. Examples of reported fusion partners include maltose-binding protein (Cardoso et al., 2015; Rahnama et al., 2017), serum albumin (Minassian et al., 2013; Neff et al., 2020), small ubiquitin-related modifier (Sermadiras et al., 2013), and antibodies (Edwards et al., 2014). Specific additional advantages of antibody fusions include: straightforward polyclonal fluorescence labelling; fragment crystallisable (Fc) chain dimerization to boost avidity; combinatorial toxin presentation to shift selectivity and/or potency; and a range of fusion/conjugation locations (Edwards et al., 2014; Wang et al., 2016; Biswas et al., 2017).

Here, we establish a straightforward strategy for the robust, high-yield production of Fc-toxin fusions. We found that monovalent Fc constituted a robust scaffold for toxin production across a variety of snake, spider and scorpion-derived peptide toxins, greatly exceeding yields versus bivalent Fc, and achieving mg quantities from only 20-70 ml cell cultures. Snake toxins retained high potency against nAChRs, with single digit nanomolar IC_50_s. Spider and scorpion toxins retained pharmacological function and selectivity across a panel of five Na_v_ subtypes. Potency, albeit reduced, often remained in the EC_50_ or IC_50_ range of 100 nM to single digit micromolar. The Fc-toxin fusions exhibited monodispersity in chromatography and remained active when kept refrigerated for over one month, while also being amenable to snap freeze-thawing for long-term storage.

## BULLET POINT SUMMARY

### What is already known?

- Previous studies provide examples of animal toxins that can be expressed recombinantly as fusion proteins.

### What does this study add?

- A methodical study on the impact of toxin valency on Fc-toxin fusion yields.
- A straightforward approach for robust recombinant production of distinct snake, spider and scorpion toxins.

### What is the clinical significance?

- Enabling toxin production empowers toxin engineering approaches to develop pharmacological tools and therapeutic leads against ion channels.

## METHODS

### Fc-Toxin Fusion Construct Design

Toxin sequences used in this study are depicted in Figure 1. For the bivalent fusions, the human immunoglobulin γ1 heavy chain wild-type protein sequence (Uniprot P0DOX5 entry 1-448 QVQL…LSPGK) was used as a template, with the following modifications. The N-terminus was truncated to begin at Lys218 in the upper hinge of the chain, so it only encoded the Fc region (218-448 KSCD…LSPGK) but retained its ability to dimerise. Three point mutations, known to attenuate Fc-dependent effector functions (L234F, L235E, and P331S), were also incorporated to generate the final bivalent Fc-toxin construct (Oganesyan et al., 2008). For the monovalent fusions, this was modified to allow for heterodimerisation through two additional, separate, knobs-into-holes mutations (Ridgway et al., 1996), which yielded two complementary constructs: Fc-knob (T366Y) and Fc-hole (Y407T). Toxins were subcloned into the pHLsec vector (Aricescu et al., 2006) between the N-terminal signal sequence and a double stop codon. Depending on downstream requirements, the N-terminal signal sequence was followed by no fusion partner and a His8 purification tag before the toxin, or bivalent Fc plus a (GGS)_12_ linker and His8 before the toxin, or monovalent Fc-knob plus a (GGS)_12_ linker and His8 before the toxin, or bivalent Fc plus a (GGS)_12_ linker and His8, followed by the toxin and a C-terminal monoVenus fluorophore (Nagai et al., 2002). The Fc-hole construct was subcloned after the signal sequence with no fusion partners or purification tags.

**FIGURE 1.**
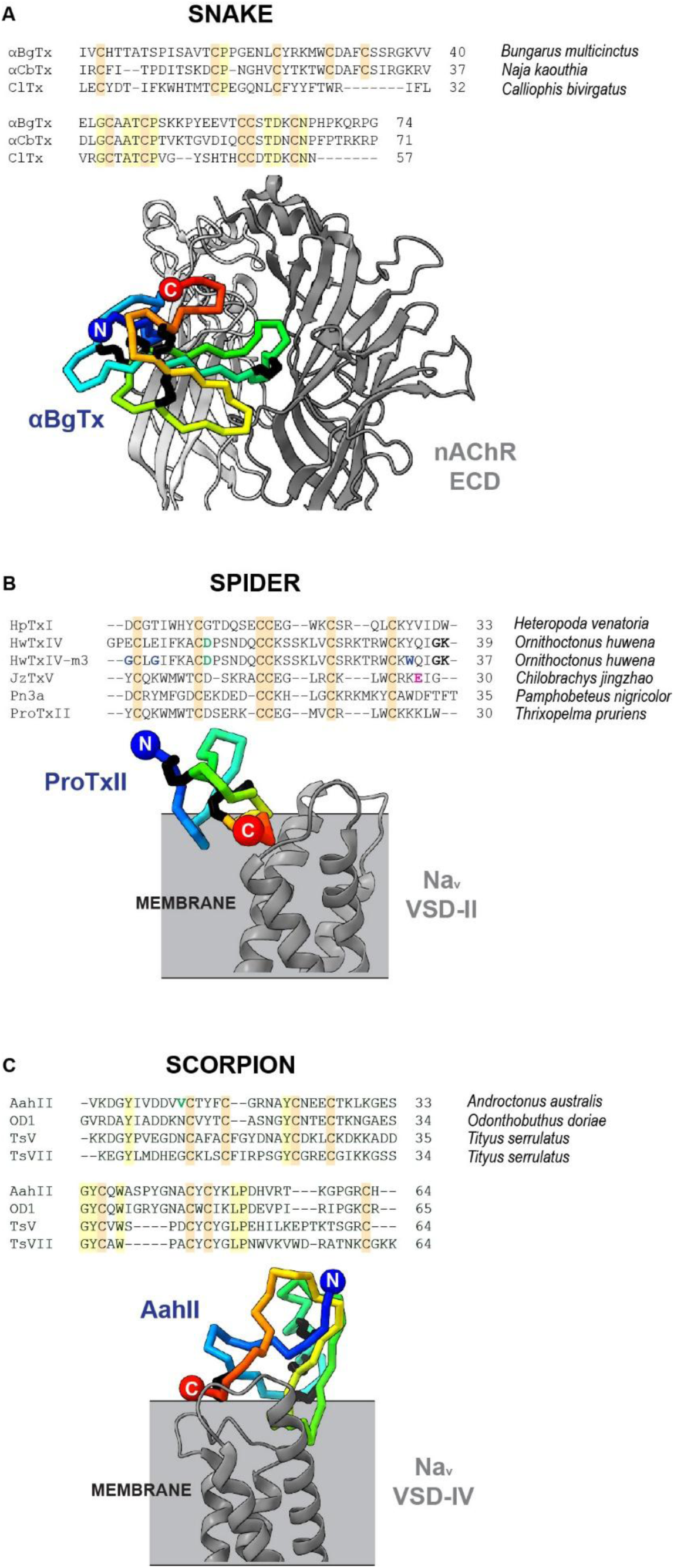
Toxin Protein Sequences and Binding Modes. **A.** Protein sequence alignment of three-finger snake toxins and structure of the *Torpedo californica* nicotinic acetylcholine receptor (nAChR) bound to αBgTx (bottom; PDB ID: 6UWZ). The extracellular domains (ECDs) of two nAChR subunits are shown, α_γ_ in grey and γ in white, with αBgTx bound across the α-γ interface. **B.** Inhibitory cystine knot spider toxin sequences and structure of chimeric voltage-gated sodium (Na_v_) channel Na_v_Ab-Na_v_1.7 bound to ProTxII (bottom; PDB ID: 6N4I). The voltage-sensing domain II (VSD-II) is shown in grey, with ProTxII bound on top, pointing diagonally away from the rest of the Na_v_. **C.** Scorpion α-toxin sequences and structure of chimeric Na_v_PaS-Na_v_1.7 channel bound to AahII (bottom; PDB ID: 6NT4). The voltage-sensing domain IV (VSD-IV) is shown in grey, with AahII bound on top, pointing diagonally away from the rest of the Na_v_. For protein sequences: cysteine residues that form disulphide bridges are shown in orange highlight; other conserved residues in yellow highlight; bold font residues represent sequence modifications to mimic C-terminal amidation (black), to increase potency (blue), to remove N-linked glycosylation sites (green), and to vary target receptor selectivity (pink). For structures: receptors are shown in greyscale ribbon format, while toxins are shown in C_α_ thick stick format with rainbow colour palettes from N-terminus (blue) to C-terminus (red), and Cys-bridges shown in black.

### Recombinant Expression, Purification & Yield Quantification

Twenty-five ml batches of Expi293F suspension cells (Thermofisher) were grown to densities of 4×10^6^ cells/ml in Expi Expression Medium (EEM, Gibco), shaking at 130 rpm, 37°C, 8% CO_2_ and transfected with 50 μg DNA and 200 μg PEI Max (Polysciences). Cells were centrifuged and resuspended in 50 ml fresh media the following day, and supernatant harvested 5 days later with cell viability typically at ∼70-80%. For bivalent/unfused expression, a plasmid encoding the bivalent Fc-toxin fusion or the unfused toxin was transfected alone. For monovalent expression, plasmids encoding the monovalent Fc-knob toxin, plus unfused Fc-hole, were co-transfected at a 1:1 ratio. Toxins were purified by addition of 5 ml binding buffer (50 mM Tris pH 7.8, 0.3 M NaCl) and 0.4 ml nickel agarose resin, gently rotated at 4°C for 1 hour. All purification buffers were supplemented with 0.02% (v/v) pluronic F-127 (Sigma-Aldrich). After washing and imidazole elution (phosphate-buffered saline (PBS) containing 20 mM Tris pH 8.0, 0.5 M NaCl, 0.25 M imidazole), final concentrations and total recombinant protein yields of the purified Fc-toxin fusions were calculated by measuring absorbance at 280 nm and correcting for extinction coefficients and elution volumes. For downstream experiments, toxins were concentrated and buffer-exchanged by ultrafiltration using 10 kDa cut-off membranes (Amicon) into extracellular electrophysiology solution (see below for composition). Average yields, for successfully expressed toxins, were ∼20 µg or ∼40 µg per ml of harvested EEM, for bivalent and monovalent fusions, respectively. Stocks were concentrated to 100 µM and kept refrigerated for up to one month before use.

### Analysis of Purity, Dimerisation, Monodispersity & Storage Conditions

To estimate the molecular mass and gauge the purity of the products obtained from recombinant expression and purification, all Fc-toxin fusions were run on SDS-PAGE. Non-reducing SDS-PAGE was used to corroborate that both bivalent and monovalent fusions remained in Fc dimer formats, while reducing SDS-PAGE was used to confirm that bivalent and monovalent fusions were composed of homodimers and heterodimers, respectively. To assess sample monodispersity and aggregation propensity, all monovalent fusions and the highest-yielding bivalent fusion were run on size-exclusion chromatography (SEC), using 70 µg loads and a Superdex 200 Increase 10/300 column (GE Healthcare) pre-equilibrated with buffer containing 300 mM NaCl, 20 mM HEPES pH 7.2, 0.02% (v/v) pluronic F-127. To investigate suitable storage conditions for all Fc-toxin fusions, a subset of the monovalent toxins was snap-frozen at 100 µM, or 5 µM, or 100 µM with a 20% (v/v) glycerol spike, and then thawed once. Samples were run on SEC alongside a non-frozen, refrigerated control to evaluate the effect of different snap freeze-thawing conditions on toxin quality, as indicated by SEC main peak size, monodispersity, and aggregation.

### Bivalent Na_v_ Toxin Function: Whole-Cell Patchliner Electrophysiology

Whole cell recordings of Na^+^ currents were made from a previously established HEK293 cell line stably expressing human Na_V_1.7 with C-terminal HAT and FLAG epitope tags (Kanellopoulos et al., 2018; Salvage et al., 2023) on a four-channel automated patch clamp system (Patchliner Quattro, Nanion Technologies). The extracellular solution contained 140 mM NaCl, 2 mM KCl, 1.5 mM CaCl_2_, 10 mM glucose, 1 mM MgCl_2_, 10 mM HEPES pH 7.4 (± 0.02 with NaOH). The intracellular solution contained 35 mM NaCl, 105 mM CsF, 10 mM EGTA, 10 mM HEPES pH 7.2 (± 0.02 with CsOH). Experiments were performed in the whole-cell configuration with HEKA EPC10 amplifiers using medium resistance (2-3 MΩ) NPC-16 chips (Nanion Technologies). Signals were low-pass Bessel filtered at a frequency of 2.9 kHz and leak currents subtracted using an online P/4 protocol. Only cells with series resistances of ≤ 8 MΩ prior to compensation (60 - 70%), seal resistances of ≥ 0.5 GΩ, and initial currents larger than 100 pA were included. Recordings commenced ≥ 5 mins from obtaining the whole cell configuration.

Cells were initially perfused 2X with normal extracellular solution and incubated for 1 min. All voltage protocols used a 50 ms duration -120 mV holding voltage. A fixed pulse protocol delivering -10 mV test pulses every 5 s for 30 pulses enabled verification of recording stability (data not shown). Control (0 µM) values were then obtained from an activation protocol delivering 100 ms test pulses ranging from -140 mV to +55 mV in 5 mV increments. Following the control period the toxins HwTxIV (10 µM) or OD1 µM (1 µM) were perfused 2X and incubated for 1 min. A series of fixed pulse protocols as described above were applied to allow for a total incubation period of approximately 8 mins (data not shown) followed by the activation protocol. Na^+^ currents were normalised against the whole-cell capacitance (C_m_) and the *I*/V relationship plotted from peak current (*I*_Na_) at each test voltage, activation curves were fitted to the Boltzmann function:

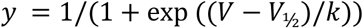

where V_½_ is the voltage of half maximal activation, *k* is the slope factor and *V* is the test voltage. Data were analysed in PatchMaster (version 2×90.5) and GraphPad Prism (version 10.3.1) with statistical comparisons made using paired t-tests.

### Monovalent Na_v_ Toxin Function: Whole-Cell Qube Electrophysiology

Automated patch clamp recordings were made from Chinese hamster ovary (CHO) cells stably expressing human Na_V_1.X α-subunits (1.1, 1.5, 1.6, 1.7, or 1.8) using the Qube384 platform (Sophion) and ViewPoint 9.0 software (Sophion). Cells were loaded onto the Qube384 in extracellular solution at a concentration of > 1 x 10^6^/ml (up to four cell lines per experiment), before positioning onto the Qchip with a pressure of -100 mbar. Whole-cell configuration was obtained using 5 x 2 s pulses of increasing pressure from -250 to -500 mbar, applied at 0.1 Hz. An adaptive voltage protocol was initially performed to determine the voltage eliciting 50% inactivation (V0.5inact) for each well, achieved by holding the cell at increasing voltages (-120 mV to 0 mV, Δ20 mV/0.03 Hz) for 8 s, prior to a 100 ms step to -120 mV, and then recording the current amplitude during a final stimulation at -10 mV (20 ms; inter-sweep voltage = -120 mV). The screening voltage protocol used for the remainder of the experiment consisted of holding the cell at V0.5inact for 8 s, before a 100 ms step to -120 mV, followed by a 20 ms stimulation at -10 mV (applied at 0.03 Hz; inter-sweep voltage = -120 mV).

Using this protocol, cells were recorded from in vehicle (0.3 % v/v DMSO) for 10 mins, then the Fc-toxin samples for 30 mins (single concentration, applied twice), and a final addition of reference inhibitor (Na_V_1.7: PF-05089771 (Tocris) (Alexandrou et al., 2016); other Na_V_1.X isoforms: tetracaine (Sigma) for 3 mins. Recordings were obtained using multi-hole acquisition at 25°C with the following recording solutions – extracellular solution: 140 mM NaCl, 11.1 mM glucose, 5 mM KCl, 5 mM HEPES, 3 mM CaCl_2_ and 1.2 mM MgCl_2_, pH 7.4, mOsm 290; intracellular solution: 120 mM CsF, 15 mM NaCl, 10 mM EGTA, 10 mM HEPES, pH 7.25, mOsm 300. All recording solutions contained (0.02% w/v) pluronic F-127, while Na_V_1.8 recording solutions also contained 0.3 µM tetrodotoxin (Tocris). Inhibition was measured at - 10 mV, as the mean current amplitude of the final three sweeps in test sample relative to the mean of the final three sweeps in vehicle. Potentiation was measured as the increase in maximal charge, during -10 mV stimulation, recorded in the test sample period relative to that of the vehicle period. Data were analysed using Analyzer (Sophion, version 9.0). Dose-response curves were constructed using GraphPad Prism (version 10.3.1) by fitting inhibition or potentiation data with a four-parameter variable slope dose-response curve, constrained between 0 – 100 % for inhibition data, and a bottom of 0 % and Hill slope = 1 for potentiation data.

### Dorsal Root Ganglion Electrophysiology

Male Wistar Han rats (21 – 27 days old) were sourced from Charles River (Cambridge, UK) and sacrificed in line with the UK Animals (Scientific Procedures) Act 1986. Dorsal root ganglia were harvested and incubated in HBSS (Gibco) containing Dispase II (Sigma, 2.5 mg/ml) for 30 min (37 °C/ 5 % CO_2_). HBSS was removed and replaced with fresh HBSS containing collagenase (Sigma, 1 mg/ml). Dorsal root ganglia were gently washed, triturated and strained, before pelleting by centrifugation (200 g, 6 min). Upon resuspension in DMEM F-12 Nutrient Mixture with Glutamax (Gibco), supplemented with 10 % (v/v) FBS, 1 % (w/v) non-essential amino acids, 50 U/ml penicillin and 50 µg/ml streptomycin, isolated neurons were seeded onto 35 mm coverslips coated with poly-D-lysine (30 µg/ml) and Laminin (Sigma, 5 µg/ml), and incubated at 37 °C/5 % CO_2_ for 1 – 2 days. Neurons were recorded in current clamp mode using an Axon MultiClamp 700B amplifier (Molecular Devices) and Clampex 11.2 software. Extracellular solution contained (in mM): 135 NaCl, 10 HEPES, 10 Glucose, 4.7 KCl, 1 MgCl_2_, 1 CaCl_2_. Intracellular solution contained (in mM): 130 KCl, 10 Glucose, 10 HEPES, 5 EGTA, 5 MgATP, 1 MgCl_2_, 0.3 NaGTP. Pipette resistance was ≤2.0 MΩ. Neurons were clamped at -70 mV and subject to an ascending ramp of current injection (500 ms, 0.1 Hz). Ramp amplitude was adjusted for each neuron to obtain an amplitude that elicits comparable multiple firing between neurons at baseline (9.2 ± 2.86 action potentials, mean ± SD). Neurons were tested in vehicle (0.01 % w/v Pluronic F-127, “baseline”), FcMoV-ProTxII (1 µM) and vehicle again (“wash-off”), with the ramp being continually applied in each condition until a repeated response was observed. Data were analysed using Clampfit11.2 (Molecular Devices) and GraphPad 10.2 (Dotmatics).

### Bivalent and Monovalent nAChR Toxin Function: Ca^2+^ Flux Assay

A modified version of the CN21 cell line, derived from TE671 cells (Merck, a human rhabdomyosarcoma cell line), which endogenously express nAChRs (Bencherif and Lukas, 1991; Beeson et al., 1996), was maintained in adherent culture at 37°C, 8% CO_2_, in high-glucose DMEM (Sigma-Aldrich) supplemented with 10% foetal calf serum (FCS, Gibco) and 1% penicillin-streptomycin-amphotericin B solution (Lonza). Sixteen hours prior to the experiment, cells were seeded at 50% confluence into flat-bottom black 96-well microplates (Corning) pre-treated with 50 µl/well of 0.05 mg/ml poly-D-lysine solution (Gibco). For the assay, 80-90% confluent cells were washed twice with buffer composed of Hank’s balanced salt solution (+Ca^2+^, +Mg^2+^, Gibco) supplemented with 20 mM HEPES pH 7.4 and incubated with 100 µl/well of loading buffer consisting of wash buffer supplemented with 2.5 mM water-soluble probenecid (Gibco), 1% (v/v) PowerLoad Concentrate (100X, Invitrogen), and 1 µM Fura-2 AM (Abcam). Cells were incubated at 37 °C, 8% CO_2_ for 45 mins and later washed twice with 100 µl/well of N-methylglucamine (NMG) buffer consisting of 0.9 mM KH_2_PO4, 0.8 mM MgSO_4_, 3 mM CaCl_2_, 25 mM glucose, 4.5 mM KCl, 130 mM N-methylglucamine, 20 mM HEPES pH 7.4. Next, 150 µl/well of toxin solution was added, comprising NMG buffer supplemented with 1.25 mM probenecid, 100 µM scopolamine hydrobromide trihydrate (Thermofisher), 0-100 nM Fc-toxin fusion, and 0.25% (w/v) bovine serum albumin (BSA). Cells were incubated at room temperature for 20 mins, before measuring fluorescence.

Fluorescence was measured with a CLARIOStar Plus microplate reader (BMG Labtech). The monochromator filter was set to read at two channels: Ca^2+^-free Ex/Em 380-12/520-30 nm, dichroic 420 nm; Ca^2+^-saturated Ex/Em 335-12/520-30 nm, dichroic 420 nm. The gain was adjusted so both channels were approximately equal and at 20% of the detector saturation intensity. Baseline signals were recorded for 10 seconds, before 50 µl 4X agonist solution, consisting of NMG buffer with 40 µM acetylcholine chloride (ThermoScientific), was injected and responses were recorded for the next 60 seconds. Fluorescence intensities (F) were measured in relative fluorescence units (RFUs). Ca^2+^ signal intensities (I) were calculated from the ratio between the fluorescence measured at the Ca^2+^-sat and Ca^2+^-free (F_sat_/F_free_). Ca^2+^ signal intensities were then normalized to their baseline signals according to the equation (I - I_0_)/I_0_, where I_0_ is the mean of the baseline Ca^2+^ signal intensities measured in the 10 second kinetic window before agonist injection. The calcium response (R) was then taken to be the maximum intensity after agonist injection (I_max_). Responses were normalized to the average responses of the positive (p) and negative (n) control wells, treated with 40 µM and 0 µM acetylcholine, respectively, according to the following equation: %R = ((R – Rn) /(Rp – Rn)) ∗ 100. Dose-response curves were fitted on GraphPad Prism (version 10.3.1), using the following 4-parameter logistic function, where C, T, B and H are the concentration, top, bottom, and Hill Slope, respectively: R(C) = B + (C^H^(T − B))/(C^H^ + EC_50_^H^).

### Mouse DRG Culturing & Staining

Adult male and female C57BL/6J mice (8–13 weeks old) were obtained from Envigo (Cambridge, UK). Mice were housed in 21°C temperature-controlled rooms with a 12-hour light-dark cycle. They were kept in groups of 4-5 in open ventilation cages and provided with nesting material, a red plastic shelter and access to chow food and drinking water *ad-libidum*. Neurons were cultured from the lumbar (L1-L5) dorsal root ganglia (DRG) of mice as previously described (Pattison et al., 2024). Briefly, animals were euthanised with rising CO_2_ and cervical dislocation. DRGs were harvested and then incubated for 15 mins at 37°C (in 5% CO_2_) in Lebovitz L-15 Glutamax (Invitrogen) media containing 1 mg/ml collagenase type 1A (Sigma-Aldrich) and 6 mg/ml BSA (Sigma-Aldrich). This was followed by a 30-mins incubation at 37°C (in 5% CO_2_) in L-15 medium containing 1 mg/ml trypsin (Sigma-Aldrich) and 6 mg/ml BSA. The DRG were gently triturated and collected by brief centrifugation (100 g). Dissociated cells in the supernatant were collected. Trituration, centrifugation and collection were repeated five times. The dissociated cells were pelleted by centrifugation at (100 g, 5 mins) and resuspended in L-15 medium containing 300 units/ml penicillin and 0.3 mg/ml streptomycin (P/S), 2.6% (v/v) NaHCO_3_, 1.5% (v/v) glucose, and 10% (v/v) FCS. The supernatant was plated onto 35 mm poly-D-lysine-coated glass bottom culture dishes (MatTek), further coated with laminin (Thermofisher), and incubated at 37°C in 5% CO_2_.

### HEK293 Culturing

HEK293 cells stably expressing Nav1.7 (Salvage et al., 2023) or HEK293T cells were maintained in high glucose Dulbecco’s modified Eagle’s medium (Sigma-Aldrich) supplemented with 10% foetal bovine serum (FBS) (Gibco), 2 mM L-glutamine (Gibco) and 100 units/mL/ 10,000 μg/mL penicillin/streptomycin (Gibco). Cells were incubated at 37°C (5% CO2, H2O) and upon reaching 80% confluency, cells were passaged using 0.05% Trypsin-EDTA (Gibco). Cells were seeded onto Poly-L-Lysine (Sigma-Aldrich)-coated 13 mm borosilicate glass coverslips (VWR) and incubated overnight to a confluency of 20-30%.

### Tertiary Cell Staining

All toxin and antibody solutions were prepared in PBS containing 0.5% BSA. Cell-coated coverslips were washed twice with 1 mL phosphate buffered saline (PBS) (Oxoid) supplemented with 0.5% bovine serum albumin (BSA) (Sigma-Aldrich) before being treated with either 100 μl FcMoV-ProTxII (10 μM) or Atto488-ProTxII (100 nM) (Montnach et al., 2021) and incubated at room temperature for 40 minutes. The cells were washed twice with 1 mL PBS/BSA (0.5%) before the addition of 100 μl Goat anti-Human IgG, Alexa Fluor™ 568 (Invitrogen, 8 µg/mL). After a 10-minute incubation at room temperature, the cells were washed twice with 1 mL PBS/BSA (0.5%) and were treated with 100 μl Donkey anti-Goat IgG, Alexa Fluor™ 568 (Invitrogen, 8 µg/mL). After a 5-minute incubation at room temperature, the cells were washed three times with 1 mL PBS and fixed with 500 μl 4% (w/v) paraformaldehyde (Sigma-Aldrich) for 10 minutes at room temperature. The coverslips were washed three-times with PBS, before mounting onto microscope slides with Aqueous DAPI mountant media (Abcam). Coverslips incubated with Atto-ProTxII were mounted directly onto microscope slides without secondary antibody incubations.

Cell images for quantification were captured less than 24 hours after staining using the EVOS M5000 imaging system at 20x magnification, Ex: 542/20, Em: 593/40 for Alexa Fluor™ 568, Ex: 482/25, Em: 524/24 for Atto 488. Fiji (Schindelin et al., 2012) was used to select 50-150 regions of interest (ROI) across a minimum of 3 images for each coverslip. 10 cell-sized selections were used to measure the average background intensity for each image and was subtracted from the mean intensity of each ROI. For representative images, the Leica Stellaris 5 was used. A HC PL APO CS2 63x/1.40 (Oil) objective lens was used, with a CS2 UV Optics 1 filter and a HyD S detector in counting mode. For all samples, a pinhole size of 95.5 µm was used, and images have a pixel size of 0.09 µm. Alexa Fluor™ 568 fluorophores were excited with a white light laser (578 nm) and light emission was detected between 584 nm – 741 nm. DAPI fluorophores were excited with a diode 405 laser (405 nm) and light emission was detected between 425 nm – 572 nm.

### Materials

Drugs/Toxins/Antibodies:

- Acetylcholine (Thermofisher)
- Tetracaine (Sigma)
- PF-05089771 (Tocris)
- Tetrodotoxin (Tocris)
- AahII (Smartox)
- OD1 (Smartox)
- HwTxIV (Alomone)
- ProTxII (Alomone)
- Atto488-ProTxII (Smartox)
- αBgTx (Invitrogen)
- αCbTx (Smartox)
- Donkey anti-Goat IgG, Alexa Fluor™ 568 (Invitrogen, A11057, AB_2534104)
- Goat anti-Human IgG, Alexa Fluor™ 568 (Invitrogen, A21090, AB_2535746)

Cell Lines:

- Expi293F Cells (Thermofisher)
- HEK293T Cells (Thermofisher)
- Modified CN21 Cell Line (Dong Lab, University of Oxford)
- Human Na_v_1.7 HEK293 Stable Cell Line (Jackson Lab, University of Cambridge)
- Human Na_V_1.1, 1.5, 1.6, 1.7 & 1.8 CHO Stable Cell Lines (Metrion Biosciences)

Details of other materials and suppliers were provided in their specific sections.

### Data Analysis & Statistics

Toxins were separately transfected and purified from 6 ml media for systematic yield quantification. Due to the large number of purifications, the cost, and the resources required, n = 3 per toxin was performed, except for the unfused FcBiV and FcMoV constructs, which were n = 6 for direct comparison by unpaired t-test. This involved > 80 total purifications (not including the larger scale 50-100 ml purifications required to prepare toxins for electrophysiology experiments), which represent larger statistical datasets. Statistical comparison per toxin was not performed because the aim was not to compare by individual toxin but to compare the FcBiV and FcMoV platforms as a whole, which had n = 39 purifications each. Statistical comparison was therefore performed by category (snake, spider, scorpion) using two-tailed Mann-Whitney U tests that allowed for analysis of non-parametric datasets with unequal variances. Toxin monodispersity by SEC to evaluate sample quality and effects of storage conditions was a preliminary study, n = 1 per toxin storage condition, to inform potential users on considerations for best maintenance of toxins. Thus, it did not require statistical analysis. Nevertheless, as for the purifications, this study was performed on 10 toxins, requiring 28 SEC runs; hence, it overall constituted a substantial dataset for informed consideration. All electrophysiological data were obtained with ≥ n = 5 separate cells. Statistical comparison of Patchliner FcBiV construct data used paired t-test, while Qube FcMoV construct data used Welch’s ANOVA test for global comparison of multiple means from parametric datasets with unequal variances, followed by post-hoc analysis with Dunnet’s T3 multiple comparisons test for individual comparisons to the control conditions. The DRG electrophysiology FcMoV-ProTxII data used one-way ANOVA test. The Ca^2+^ flux assay for snake toxins acting at nAChRs was limited to n = 4 separate experiments due to limited sample stocks of valuable synthetic and purified proteins, and statistical analysis was therefore not performed. Statistical comparison of the fluorescence quantification data was performed using an unpaired t-test.

## RESULTS

### Toxin Information & Rationale for Fc-Toxin Fusion Construct Design

We selected a broad panel of 13 snake, spider, and scorpion venom-derived peptide toxins (Table 1). The panel included three snake toxins (Figure 1A), six spider toxins (Figure 1B), and four scorpion toxins (Figure 1C), each individually distinct in sequence and known for modulation of two important pharmacological and therapeutic targets, nAChRs or Na_v_s. Human codon-optimised synthetic DNAs encoding native toxin sequences were used for this study, but with modifications in some instances to mimic amidation at the C-termini (HwTxIV, HwTxIV-m3), to enhance potency (HwTxIV-m3), to modify selectivity (JzTxV), or to remove N-linked glycan sites (AahII, HwTxIV, HwTxIV-m3) (Figure 1).

A preliminary expression trial of the 13 unfused toxins from transiently transfected Expi293F cells revealed detectable toxin protein bands secreted in the media for only three of 13 toxins: 2/3 snake toxins, 0/6 spider toxins, and 1/4 scorpion toxins (Figure S1A-C). We therefore decided to investigate toxin fusion to a bivalent Fc (FcBiV) scaffold due to its high stability and expression (Lo et al., 1998). Furthermore, bivalency might be able to boost affinity through avidity effects, thus enhancing the toxin potency for its target.

High-resolution structures for each toxin class bound to its receptor target, snake toxin αBgTx-nAChR (Rahman et al., 2020), spider toxin ProTxII-Na_v_ (Xu et al., 2019), and scorpion toxin AahII-Na_V_ (Clairfeuille et al., 2019), reveal that in all cases, the N-terminus is located farther away from the receptor contact zone than the C-terminus and does not participate directly in toxin binding (Figure 1). Thus, although different to the situation in nature where the Fc is attached to the C-terminus of the antibody variable domains that target antigens, in this case, to minimise risk of steric hindrance to function, the Fc was fused to the toxin N-terminus. Furthermore, RNA translation of the Fc domain before the toxin domain might help to stabilise toxin folding, hence boosting FcBiV toxin expression.

### Expression, Purification, Yield Quantification & Function of Bivalent Fc-Toxins

The FcBiV toxins were recombinantly expressed from Expi293F cells (Figure 2A-B) and the resultant media run on reducing SDS-PAGE to visualise Fc-toxin monomers. Protein bands were not visible for AahII and TsVII, but in all other cases were visible, although only four toxins, HpTxI, HwTxIV, HwTxIV-m3, and OD1, exhibited bands visibly stronger than that of endogenously secreted human serum albumin (HSA) (Figure 2C). This represented an improvement over the benchmark expression trial of unfused toxins, which only yielded visible bands for αBgTx, αCbTx, and OD1 (Figure S1).

**FIGURE 2.**
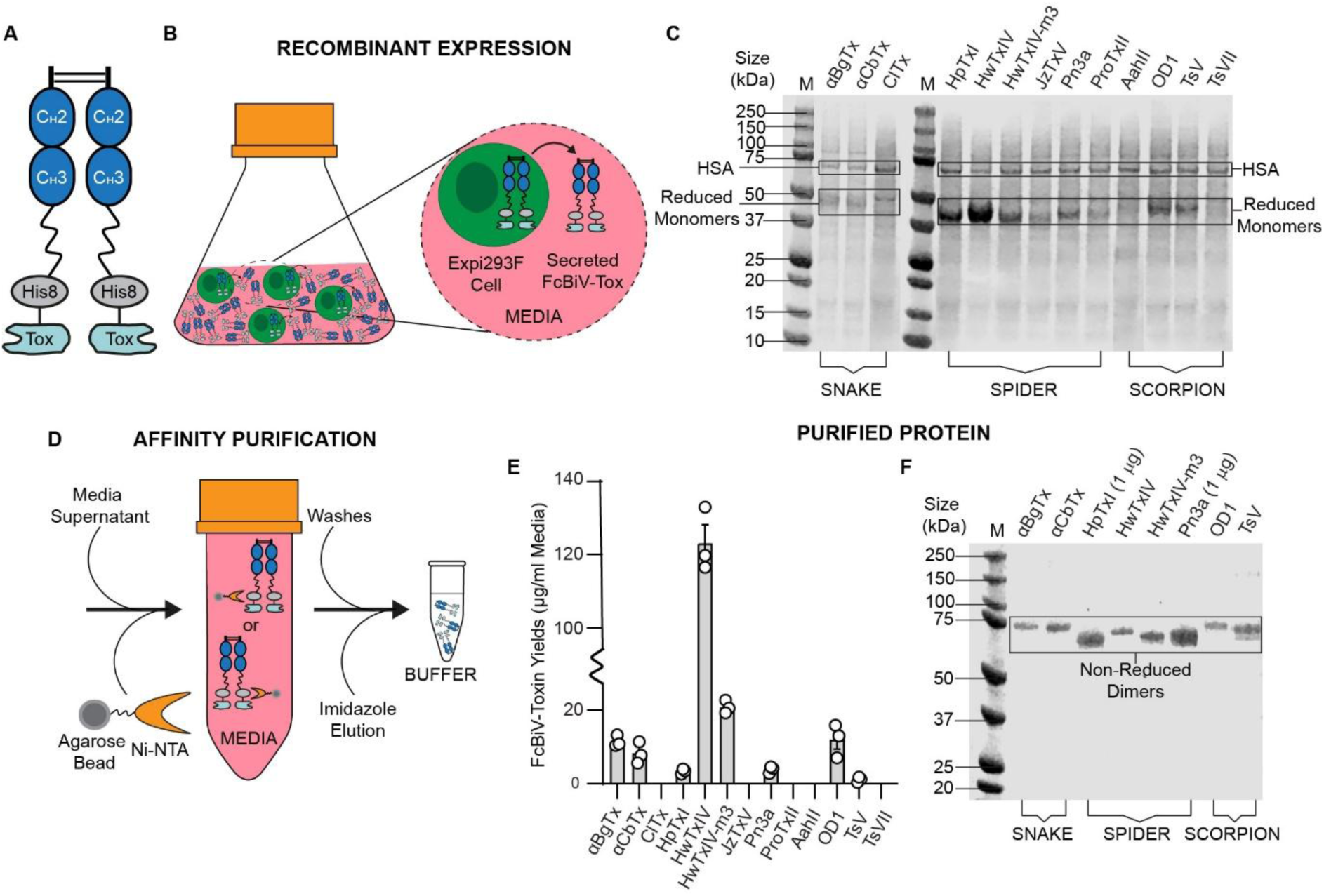
Expression, Purification & Yield Quantification of Bivalent Fc-Toxins. **A.** Schematic of a bivalent fragment crystallisable (Fc)-toxin fusion, showing its homodimeric Fc chain arrangement with two toxin copies, one at the C-terminus of each chain. **B.** Schematic of recombinant expression of bivalent Fc-toxin (FcBiV) fusions from Expi293F cells. **C.** Coomassie stained reducing SDS-PAGE gel of Expi293F expression media supernatant (cells removed; 10 μl loads), showing FcBiV toxin expression levels (bands marked “reduced monomers”). **D.** Schematic of affinity purification of FcBiV toxins from media using Ni-NTA agarose beads. **E.** Purified FcBiV toxin yields, µg protein per ml of expression media (n = 3 per toxin; n = 39 total data points). **F.** Coomassie stained non-reducing SDS-PAGE gel of the eight purified FcBiV toxins that gave detectable post-purification yields, illustrating high purity and intact homodimerisation states. 0.3 - 1 μg protein was loaded per well; band intensities are not indicative of protein yields. C_H_2/C_H_3 = constant heavy chain region 2/3; His8 = octaHis purification tag; M = size marker; HSA = human serum albumin.

Protein was affinity-purified from the media via an octaHis tag using Ni-NTA agarose beads and yields were optically evaluated after purification, revealing large variation across the panel (Figure 2D-E). From thirteen toxins, five were undetectable. For the remaining eight, with the exception of the two FcBiV HwTxIV-variant spider toxins, which both exceeded yields of 20 μg from 1 ml media (n = 3, across three separate transfections and purifications, n = 6 total results), the other FcBiV toxins yielded at or below 12 µg from 1 ml media (n = 3, across three separate transfections and purifications, n = 39 total results). The eight toxins giving detectable yields were all highly pure when run as intact dimers on non-reducing SDS-PAGE (Figure 2F). At yields of 12 μg or less from 1 ml media, 80 ml or more of expression media is required to purify 1 mg of Fc-toxin fusion. This is expensive and sub-optimal compared to typical antibody yields, which are often 8 mg or greater from 80 ml media (Fang et al., 2017).

To evaluate function of the two FcBiV snake toxins, we tested inhibition of acetylcholine evoked dose-dependent Ca^2+^ mobilisation in modified CN21 cells by inhibiting endogenously expressed nAChRs. Both αBgTx and αCbTx exhibited strong inhibition of nAChR-evoked Ca^2+^ mobilisation, achieving single digit nanomolar potency and remaining within 2-fold of synthetic toxin potency (Figure 3A-B). To evaluate function of the best-expressing spider and scorpion toxins, HwTxIV and OD1, respectively, we tested them using whole-cell patch clamp electrophysiology. Applying HwTxIV (10 µM) or OD1 (1 µM) to HEK293 cells stably expressing human Na_v_1.7, revealed that both FcBiV toxins were functional, showing statistically significant inhibition or potentiation, respectively, of Na_v_1.7 currents (Figure 3C-H). HwTxIV resulted in a 45% decrease of peak *I*_Na_ (-99.92 ± 14.06 pA/pF vs -54.57 ± 11.84 pA/pF, *n* = 6, *P* = 0.0012) without effect on the V½ of activation (*P* = 0.852, *n* = 6), consistent with previous reports (Xiao et al., 2008). OD1 resulted in a 40% increase of peak *I*_Na_ (-83.48 ± 13.28 pA/pF vs -117.67 ± 25.42 pA/pF, *n* = 6, *P* = 0.0477) and ∼5.5 mV leftward shift of V½ activation (-15.18 ± 2.32 mV vs -20.67 ± 2.88 mV, *n* = 6, *P* = 0.0178), again consistent with previous reports (Maertens et al., 2006; Salvage et al., 2023).

**FIGURE 3.**
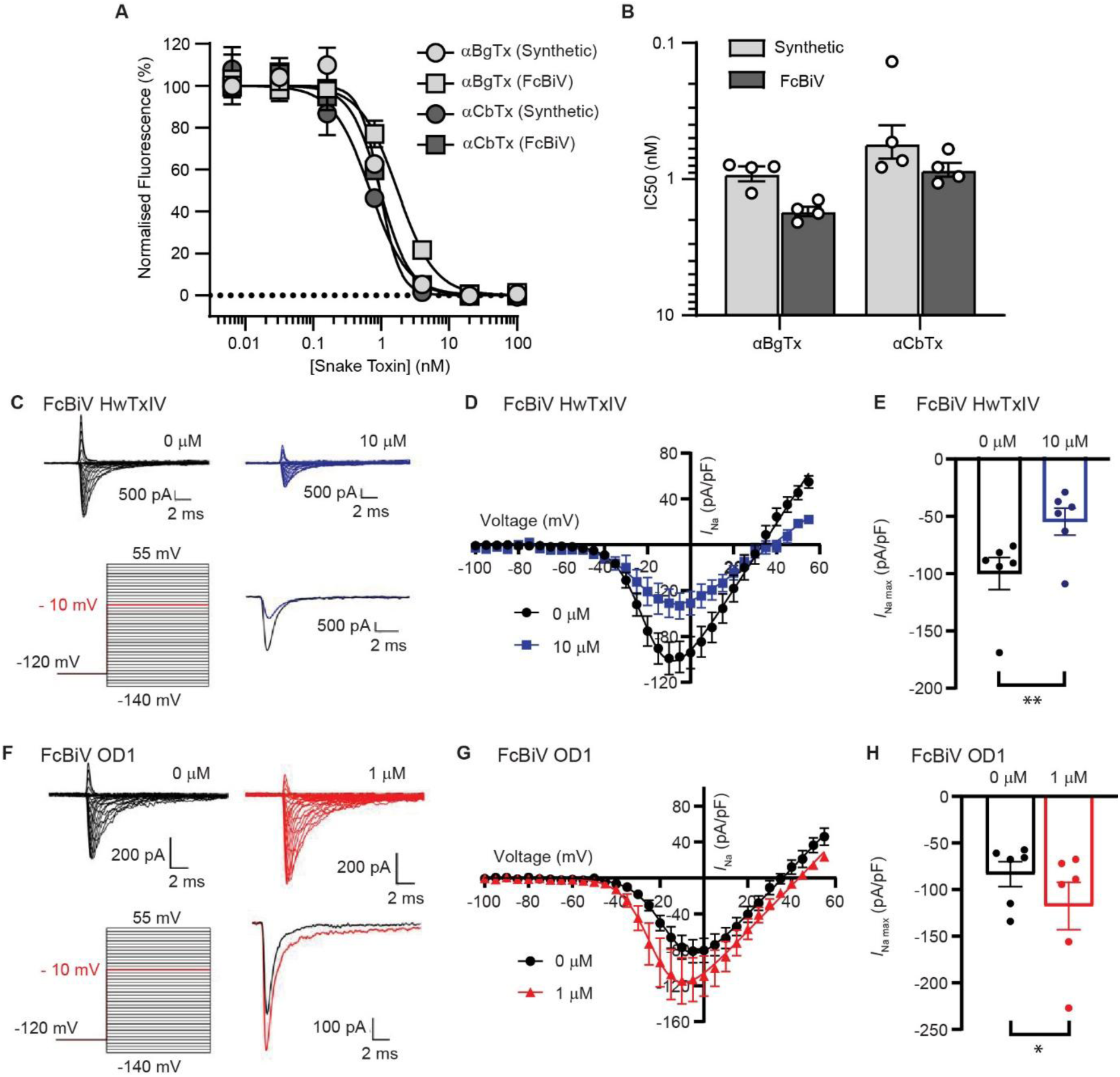
Function of Bivalent Fc-Toxins. Two FcBiV snake toxins, αBgTx and αCbTx, were functionally evaluated for inhibition of nAChR-evoked Ca^2+^ mobilisation in CN21 cells. The best-expressing FcBiV spider and scorpion toxins, HwTxIV and OD1, respectively, were functionally evaluated with whole-cell patch clamp electrophysiology for inhibition or potentiation, respectively, of Na^+^ currents in HEK293T cells stably expressing Na_v_1.7. **A.** Dose-dependent inhibition by synthetic and FcBiV αBgTx and αCBTx snake toxins on fluorescent signal generated from the action of 10 μM acetylcholine on nAChR-evoked Ca^2+^ flux in CN21 cells. **B.** Bar chart showing potencies of synthetic versus FcBiV snake toxins (n = 4 per toxin), expressed as IC_50_s. **C.** Representative traces of FcBiV HwTxIV at 10 µM with paired 0 µM control period, and voltage steps – red step corresponds to the single trace comparison. **D.** Corresponding *I*V relationships (mean ± SEM). **E.** Corresponding peak *I*_Na_ from the IV curves. **F.** Representative traces of FcBiV OD1 at 1 µM with paired 0 µM control period, and voltage steps – red step corresponds to the single trace comparison. **G.** Corresponding *I*V relationships (mean ± SEM). **H.** Corresponding peak *I*_Na_ from the IV curves.

### Expression, Purification and Yield Quantification of Monovalent Fc-Toxins

An alternative Fc scaffold setup involves using Fc heterodimers fused to a single toxin to make a monovalent binder. For this, Fc domains are engineered using knobs-into-holes mutations, such that an Fc chain with a knob or hole mutation cannot homodimerize, and instead heterodimers form between complementary Fc-hole and Fc-knob chains (Ridgway et al., 1996) (Figure 4A). We hypothesised that reducing the peptide toxin burden on the Fc dimer by reducing the toxin copy number from two to one might increase the yield for FcMoV toxin versus FcBiV toxin constructs. We first verified that the FcMoV format itself does not unduly impact expression and purification yield versus the FcBiV format. Unfused FcMoV and FcBiV formats both yielded ∼150 µg purified protein from 1 ml media (Figure S2A-B). Next, FcMoV toxins were recombinantly expressed from Expi293F cells (Figure 4B) and the resultant media run on reducing SDS-PAGE to visualise Fc-toxin monomers. In contrast to FcBiV toxins (Figure 2C), FcMoV toxins gave strong expression bands in 12/13 cases, being the predominant protein bands, visibly exceeding the HSA band (Figure 4C). Protein purification (Figure 4D) revealed that only 2 toxins, ClTx and TsVII failed to give detectable yields, whereas the remaining 11 FcMoV toxins, all gave yields at or above 15 μg from 1 ml media (n = 3, across three separate transfections and purifications per toxin), mean = 41 ug from n = 33 total results (Figure 4E). Visualisation on non-reducing SDS-PAGE revealed uniformly high purity and intact dimerisation states (Figure 4F). For all three categories, snake toxins (n = 6), spider toxins (n = 18) and scorpion toxins (n = 9), mean yields were significantly higher for FcMoV versus FcBiV toxins, by 4-fold, 2-fold, and 6-fold, respectively (Figure 4G). More importantly, six toxins from eleven that previously gave no yield (JzTxV, ProTxII and AahII) or negligible yield of 1-3 μg from 1 ml media (HpTxI, TsV, Pn3a) could now be purified at yields greater than 15 μg per ml media for downstream applications.

**FIGURE 4.**
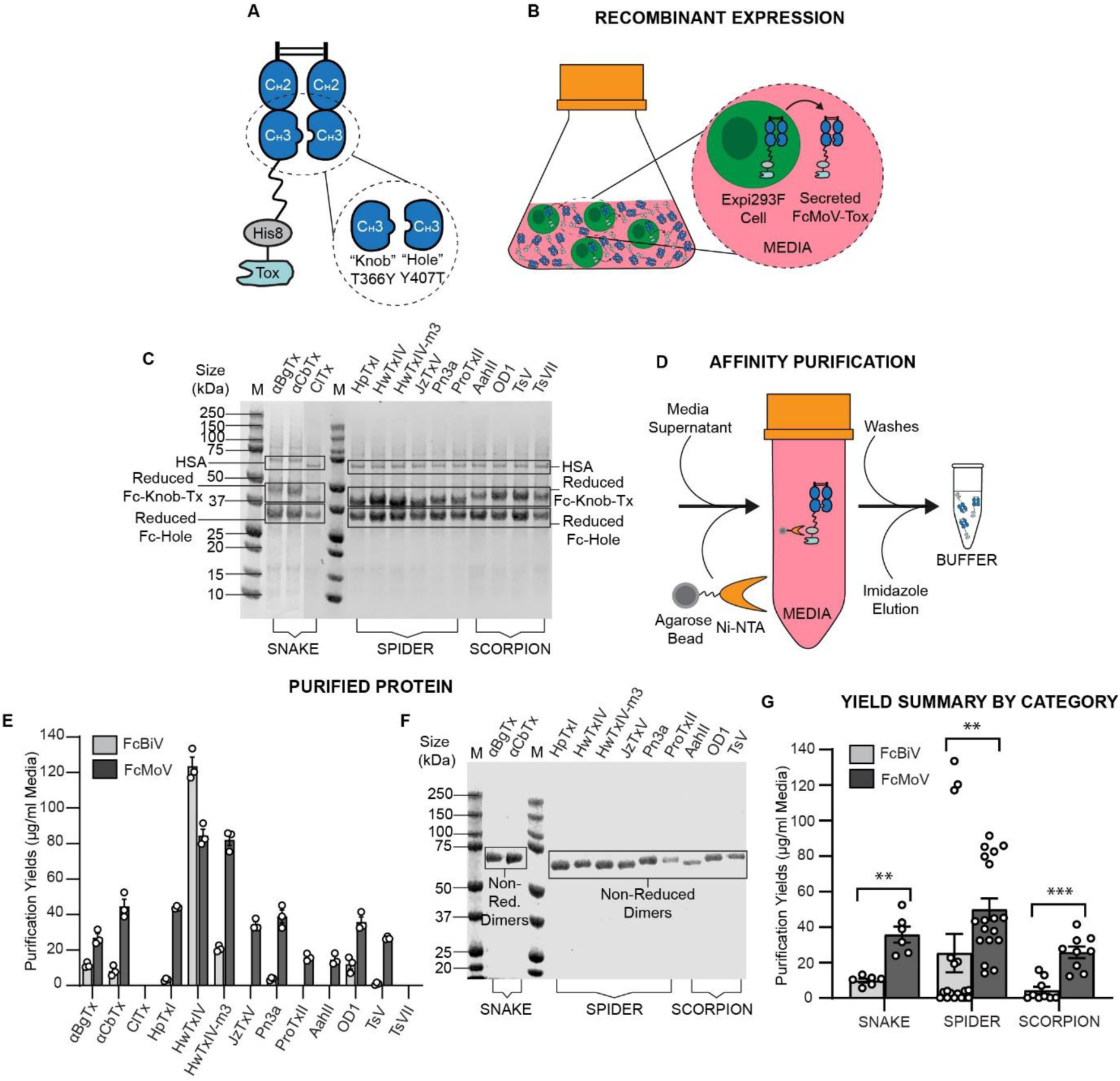
Expression, Purification & Yield Quantification of Monovalent Fc-Toxins. **A.** Schematic of a monovalent Fc-toxin fusion showing one toxin copy attached to the Fc-knob chain C-terminus and no toxin on the Fc-hole chain. **B.** Schematic of recombinant expression of monovalent Fc-toxin (FcMoV) fusions from Expi293F cells. **C.** Coomassie stained reducing SDS-PAGE gel of Expi293F expression media supernatant (cells removed; 10 μl loads). **D.** Schematic of affinity purification of FcMoV toxins from media using Ni-NTA agarose beads. **E.** Bar chart showing and comparing FcBiV versus FcMoV toxin yields, µg per ml of expression media (n = 3 per toxin; n = 78 total data points). **F.** Coomassie stained non-reducing SDS-PAGE gel of purified FcMoV toxins with detectable post-purification yields. 0.3 - 1 μg protein was loaded per well; band intensities are not indicative of protein yields. **G.** Bar chat presenting statistical comparison of FcBiV versus FcMoV yields by toxin category (snake (3-finger), spider (inhibitory cystine knot) or scorpion (α-toxin)) (n = 6-18; Mann-Whitney U tests, two-tailed; ** = p < 0.01; *** = p < 0.001). C_H_2/C_H_3 = constant heavy chain region 2/3; His8 = octaHis purification tag; M = size marker; HSA = human serum albumin.

To fully challenge the robustness of this expression format, we tested four ant venom-derived toxins: Pc1a, Ta3a, Rm4a, and Mri1a (Robinson et al., 2023). These represent a different toxin category, lacking disulphide bridges and instead being highly hydrophobic (Kyte-Doolittle scores = 0.96, 0.90, 0.70, and 1.04, respectively) (Kyte and Doolittle, 1982) (Figure S3A). Fusion of the Fc to these toxins, including trialling three different Fc set-ups for Ta3a (the peptide with the lowest number of hydrophobic residues), resulted in no detectable expression regardless of toxin valency (n = 6 total toxins in this category) (Figure S3B).

### Fc-Toxin Monodispersity and Storage

We performed a preliminary study to evaluate sample monodispersity and aggregation propensity at n = 1 per toxin condition tested. Overall, this represents a dataset of 28 toxin conditions for informed consideration of protein quality and optimal storage conditions. We assessed the monodispersity of all the purified FcMoV toxins, and the highest-yielding HwTxIV FcBiV toxin, by size-exclusion chromatography (SEC) using 70 ug loads. The snake toxins, αBgTx and αCbTx (Figure 5A-B, S4A), showed monodisperse ∼70 kDa single peaks after being snap freeze-thawed at 100 µM, hence exhibiting suitability for long-term storage at high concentrations. The scorpion toxins, AahII, OD1, and TsV, all showed monodisperse ∼70 kDa single peaks when stored on at 0°C for one month at 100 μM but accumulated an aggregation and/or a pre-shoulder peak after snap freeze-thawing at 100 μM (Figure 5C-D, S4B-C).

**FIGURE 5.**
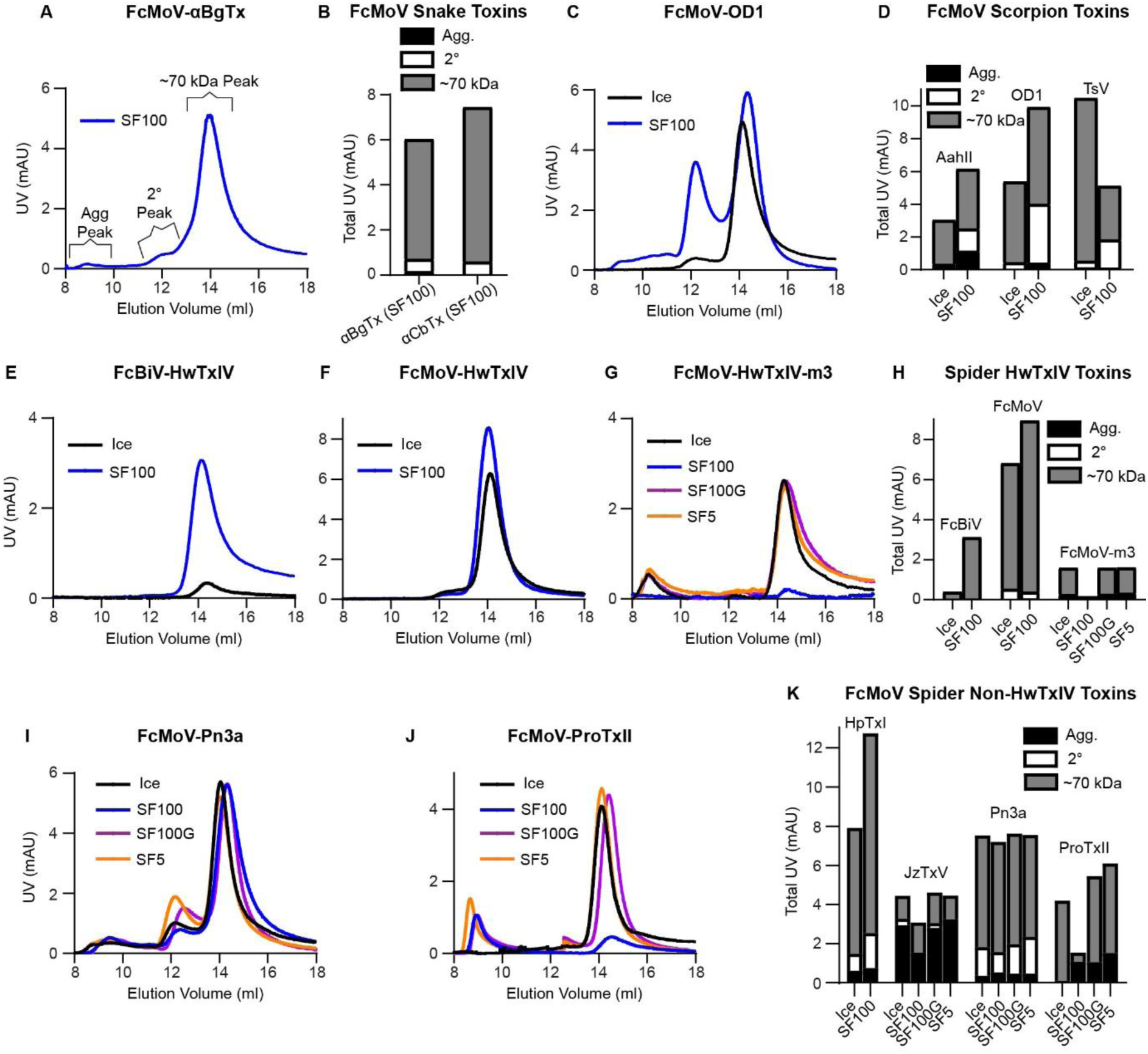
Fc-Toxin Monodispersity & Storage. All purified FcMoV toxins, and the highest-yielding purified FcBiV toxin (HwTxIV), were run on size-exclusion chromatography (SEC) using 70 µg loads to explore protein monodispersity and optimal storage conditions (n = 1 run per toxin condition; n = 28 total SEC runs). For SEC traces, ultraviolet (UV) absorbance peaks at 14-15 ml elution volume represented the ∼70 kDa monomeric Fc toxin species size, indicative of high protein quality. A higher order aggregation (agg) peak at 8-10 ml elution volume, or secondary (2⁰) shoulder peak at 11-13 ml elution volume were sometimes also visible. SEC trace colours: black (Ice) = stored on ice; blue (SF100) = snap-freeze thawed at 100 μM; purple (SF100G) = snap freeze-thawed at 100 µM with 20% glycerol; orange (SF5) = snap freeze-thawed at 5 µM. For summary bar charts, each bar represents total UV absorbance across an SEC trace from the ∼70 kDa peak (grey bar), 2⁰ peak (white bar), and agg peak (black bar). **A.** FcMoV-αBgTx trace. **B**. FcMoV snake toxin bar chart summary. **C.** FcMoV-OD1 traces. **D.** FcMoV scorpion toxin bar chart summary. **E.** FcBiV-HwTxIV traces. **F.** FcMoV-HwTxIV traces. **G.** FcMoV-HwTxIV-m3 traces. **H.** HwTxIV bar chart summary. **I.** FcMoV-Pn3a traces. **J.** FcMoV-ProTxII traces. **K.** FcMoV spider “other” toxin bar chart summary.

The spider toxins exhibited variable SEC behaviour. For HwTxIV, the FcBiV format precipitated when stored at 0°C within one or two weeks but could be preserved by snap-freezing (Figure 5E, 5H), whereas the FcMoV format remained stable when stored at 0°C and after snap freezing-thawing at 100 μM (Figure 5F, 5H). This supports the hypothesis that reducing toxin load on the Fc from two copies to one reduces aggregation propensity and increases monodisperse stability. HwTxIV-m3 also remained monodisperse after one month storage at 0°C; however, the sample was no longer detectable by SEC after snap freeze-thawing at 100 μM. Despite this, sample loss was completely rescued by adding 20 % glycerol to the 100 μM stock or reducing the concentration to 5 μM before snap-freezing (Figure 5G-H). For HpTxI and Pn3a, both were predominantly monodisperse under all conditions tested, albeit with a minor pre-shoulder (Figure 5I, 5K, S4D). In contrast, the only toxin to exhibit a significant aggregation peak under all conditions was JzTxV (Figure 5K, S4E). ProTxII behaved similarly to HwTxIV-m3, being monodisperse after storage at 0°C for one month but undetectable after snap-freezing at 100 μM, and this was rescued by adding 20 % glycerol to the 100 μM stock, or reducing the concentration to 5 μM, before snap-freezing (Figure 5J-K). For comparison, unfused FcBiV and FcMoV constructs, snap freeze-thawed at 100 μM, remained monodisperse on SEC, and serve as markers of protein peak height and elution volume (Figure S4F-G).

### Function of Monovalent Fc-Toxins

As with the FcBiV snake toxins, we tested the two FcMoV snake toxins in the Ca^2+^ flux assay. Both αBgTx and αCbTx exhibited strong inhibition of nAChR-mediated Ca^2+^ mobilisation, achieving single digit nanomolar potency that remained within 5-fold of the synthetic toxins (Figure 6A-B) and 2-fold of the FcBiV snake toxins (Figure 3A-B), suggesting the FcBiV constructs were not markedly benefitting from avidity effects to boost potency.

**FIGURE 6.**
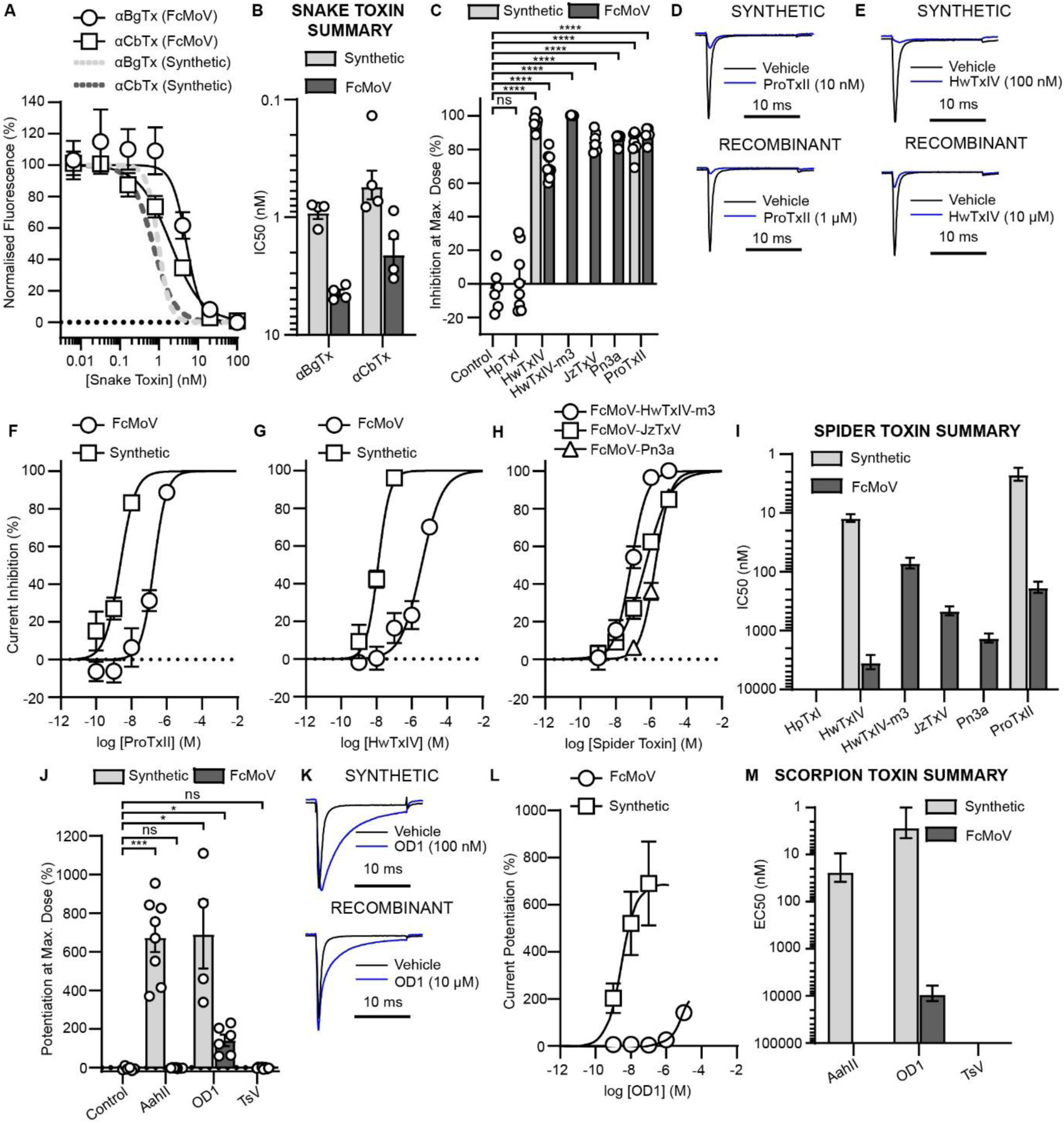
Function of Monovalent Fc-Toxins. **A.** Dose-dependent inhibition by synthetic and FcMoV αBgTx and αCBTx snake toxins on fluorescent signal generated from the action of 10 μM acetylcholine on nAChR-evoked Ca^2+^ flux in CN21 cells. **B.** Bar chart showing potencies of synthetic versus FcMoV snake toxins (n = 4 per toxin), expressed as IC_50_s **C.** Bar chart showing synthetic and FcMoV spider toxin inhibition at highest dose tested (1-10 μM) versus a negative control protein (a non-modulatory VHH) as measured using whole-cell patch clamp recordings of Na^+^ currents in CHO cells stably expressing Na_V_1.7. **D.** Representative current traces in the absence (black trace) and presence (blue trace) of synthetic ProTxII or recombinant FcMoV-ProTxII. **E**. Same as **D**, but for HwTxIV. **F.** Dose-dependent inhibition by synthetic ProTxII and FcMoV-ProTxII. **G.** Same as **F**, but for HwTxIV. **H.** Dose-dependent inhibition profiles for FcMoV fusions with HwTxIV-m3, JzTxV, and Pn3a. **I.** Bar chart showing relative potencies of synthetic versus FcMoV spider toxins, expressed as IC_50_s. **J.** Bar chart showing synthetic and FcMoV scorpion toxin potentiation at highest dose tested (100 nM for synthetic; 10 μM for FcMoV fusions) versus a negative control protein (a non-modulatory VHH). **K.** Representative current traces for synthetic (top) and FcMoV (bottom) OD1. **L.** Dose-dependent potentiation by synthetic and FcMoV-OD1. **M.** Bar chart showing relative potencies of synthetic versus FcMoV scorpion toxins, expressed as EC_50_s. All electrophysiology data points are means of n = 5-8 individual cells tested per dose. Significance of inhibition determined using Welch’s one-way ANOVA test plus post-hoc Dunnet’s T3 multiple comparisons test; ns = not significant; * = p < 0.05; *** = p < 0.001; **** = p < 0.0001).

To evaluate function of the spider and scorpion toxins, we performed whole-cell patch clamp electrophysiology using the automated high-throughput Qube platform. The long incubation times required for toxins to approach equilibrium of effect (up to 30 minutes) precluded reliable recordings of more than one concentration per cell. Therefore, Qube recordings allowed for a single concentration per cell (∼30 minutes each) from a total of ∼900 cells and data points across potency and selectivity experiments. It could also independently verify the inactivation profile per cell and set an appropriate 50% inactivation voltage individually tailored to each cell, thus accounting for variability to ensure fair comparison of toxin effects across all tested cells, both for inhibition of Na_v_ activation (spider toxins) and for inhibition of Na_v_ fast inactivation resulting in potentiation (scorpion toxins) (Figure S5A) (Bosmans and Swartz, 2010; Ahmadi et al., 2020).

For evaluation we used a CHO cell line stably expressing human Na_v_1.7, a target of interest for the treatment of chronic pain (Eagles et al., 2020). We first evaluated the toxins both for percentage modulation and sensitivity. Five of the six FcMoV spider toxins produced significant inhibition versus the no toxin control, ranging from 70-100 % at the highest dose tested (1-10 μM) (Figure 6C-E). Only FcMoV-HpTxI failed to inhibit Na^+^-mediated currents (Figure S5B). The five active toxins retained IC_50_s in the nanomolar to single digit micromolar range despite being produced recombinantly from mammalian cells and being fused to Fc domains, although potency was reduced by two to four orders of magnitude versus synthetic counterparts. ProTxII and HwTxIV IC_50_s were 190 ± 40 nM and 3550 ± 970 nM, respectively, representing 80-fold and 290-fold losses of potency relative to synthetic counterparts (Figure 6F-G, 6I). HwTxIV-m3 (70 ± 10 nM), JzTxV (468 ± 80 nM), and Pn3a (1710 ± 230 nM) IC_50_s represented reductions in potency of 180-fold (Revell et al., 2013), 740-fold (Moyer et al., 2018) and 2,140-fold (Deuis et al., 2017), respectively, versus published values (Figure 6H-I). For the three FcMoV scorpion toxins, OD1 achieved significant potentiation of ∼140 % at the highest dose tested (10 μM), albeit with potency reduced 3,470-fold relative to its synthetic counterpart (Figure 6J-M). Recombinant TsV and AahII failed to elicit significant modulation (Figure 6J, 6M, S5C-D).

We subsequently evaluated durability of activity by testing the four most effective spider toxins, HwTxIV-m3, JzTxV, Pn3a and ProTxII, when prepared fresh versus after being stored at 0°C for one month. Potency is retained within 3-fold during this timeframe (Figure 7A-D). We also tested the selectivity profiles of these four toxins by measuring inhibition against five human Na_v_ isoforms stably expressed in CHO cells: Na_v_1.1, Na_v_1.5, Na_v_1.6, Na_v_1.7, and Na_v_1.8. Consistent with previous reports for synthetic toxins, all four FcMoV toxin fusions were most potent on Na_v_1.7 (Table 2). HwTxIV-m3 exhibited 6-fold selectivity for Na_v_1.7 over Na_v_1.1 and Na_v_1.6 and was completely selective versus Na_v_1.5 or Na_v_1.8 (Figure 7E). Up to concentrations of 10 µM, JzTxV and Pn3a were completely selective for Na_v_1.7 over the other isoforms (Figure 7F-G). ProTxII exhibited no inhibition of Na_v_1.5, and IC_50_s against Na_v_1.1, Na_v_1.6 and Na_v_1.8 increased 30-fold, 90-fold, and 30-fold, respectively, such that at 1 μM, ProTxII achieved near-full inhibition of Na_v_1.7 versus almost no inhibition of other tested isoforms (Figure 7H). To demonstrate that the recombinantly produced FcMoV-toxins can be used to modulate physiological neuronal excitability, we tested ProTxII on dissociated DRG cultures, which express Na_V_1.7 channels along with other Na_V_ subtypes (Zeisel et al., 2018). Na_V_1.7 has a key role in controlling subthreshold amplification and action potential initiation in rodent and human DRG neurons (Rush et al., 2007; Alexandrou et al., 2016). Application of ProTxII (1 µM) led to a significant reduction in action potential firing without a reduction in amplitude, consistent with selective inhibition of Na_V_1.7 (Figure 7I-K).

**FIGURE 7.**
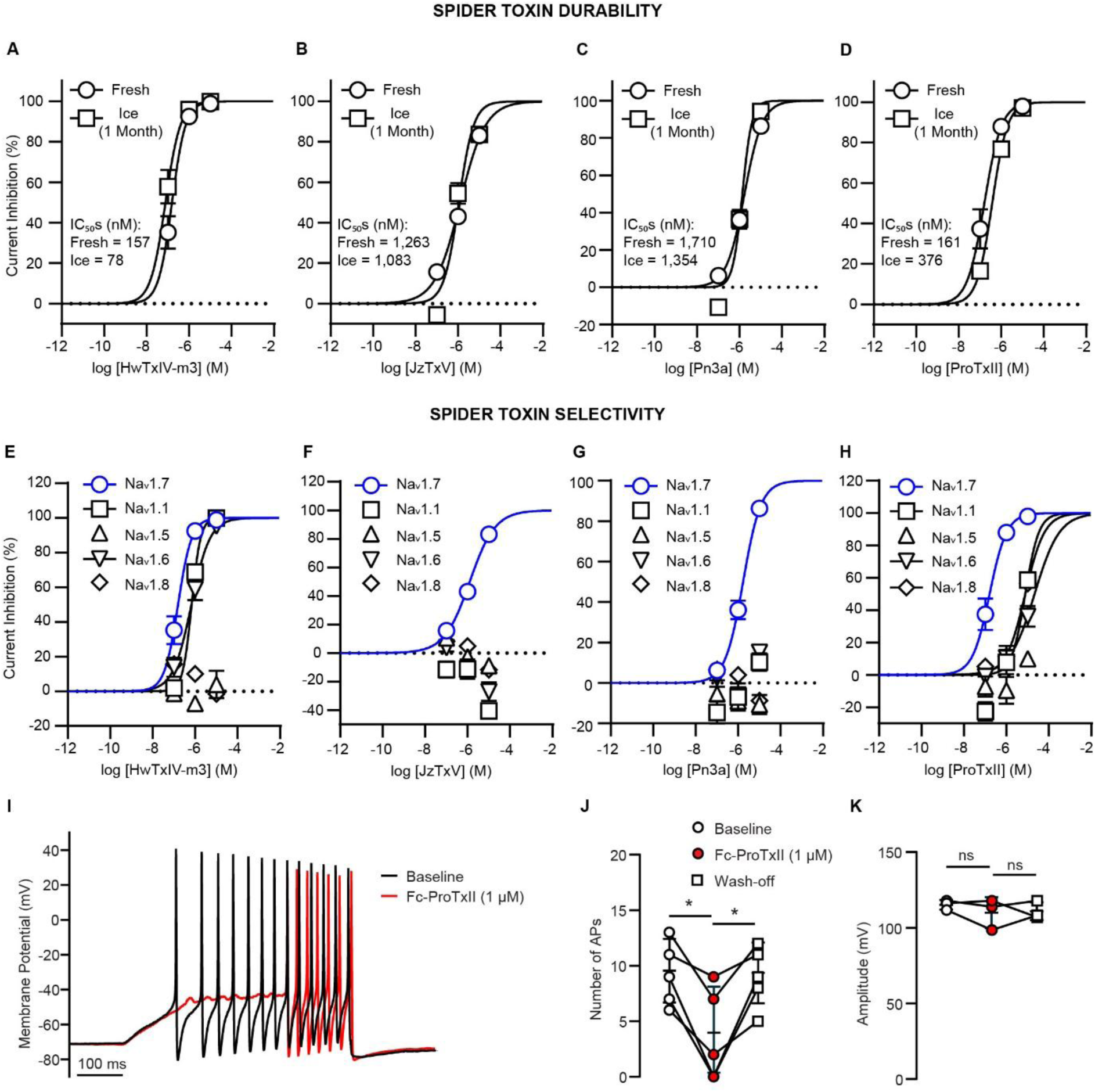
Durability & Selectivity of Monovalent Spider Fc-Toxins. Dose response profiles were measured using whole-cell patch clamp recordings of Na^+^ currents in CHO cells stably expressing Na_V_ isotypes. **A.** FcMoV-HwTxIV-m3 potency when freshly prepared versus stored on ice for one month against Na_v_1.7. **B.** Same but for FcMoV-HwTxIV-m3. **C.** Same but for FcMoV-Pn3a. **D.** Same but for FcMoV-ProTxII. **E.** FcMoV-HwTxIV-m3 potency comparison against Na_v_1.1, Na_v_1.5, Na_v_1.6, Na_v_1.7, and Na_v_1.8. **F.** Same but for FcMoV-HwTxIV-m3. **G.** Same but for FcMoV-Pn3a. **H.** Same but for FcMoV-ProTxII. All electrophysiology data points are means of n = 5-8 individual cells tested per dose. **I.** Representative current clamp traces demonstrating reversible inhibition of DRG action potential firing by FcMoV-ProTxII (1 µM). Action potentials were evoked by an ascending ramp, set at an amplitude capable of eliciting comparable multiple firing between neurons. **J.** Number of action potentials was significantly reduced following application of ProTxII in all neurons tested (n = 5). Effects of ProTxII were reversible upon wash-out. **K.** Action potential amplitude was not significantly reduced in neurons that retained firing in the presence of ProTxII (n = 3).

### Monovalent Fc-ProTxII Staining of HEK Cells and Dorsal Root Ganglia Neurons

Given the ability of the recombinantly produced Fc-toxin fusions to modulate ion channel function we also investigated these reagents for direct visualisation of targets. For this we chose FcMoV-ProTxII against Na_v_1.7, which could be benchmarked against synthetic fluorescent ATTO488-ProTxII, recently developed to address the shortage of good antibodies for visualisation of this subtype (Montnach et al., 2021). Staining with FcMoV-ProTxII plus a fluorescent secondary Alexa Fluor^TM^ (AF) 568 antibody versus ATTO488-ProTxII on the HEK293T Na_v_1.7 stable cell line or naïve HEK293T cells revealed no specific detectable signal from either probe (Figure 8A-B). Notably, the presence of the Fc domain on the FcMoV-ProTxII means the toxin signal can be amplified through polyclonal binding. To take advantage of this we introduced an additional tertiary AF568 antibody binding step (see Methods). This combination resulted in a significantly higher specific fluorescence against the HEK293T Na_v_1.7 stable cell line than naïve HEK cells, observable as surface distributed halos by confocal imaging (Figure 8C-E). We subsequently applied this strategy to staining of dissociated DRG neurons and again observed higher fluorescence versus a negative control of secondary/tertiary antibody mix alone (Figure 8F-G).

**FIGURE 8.**
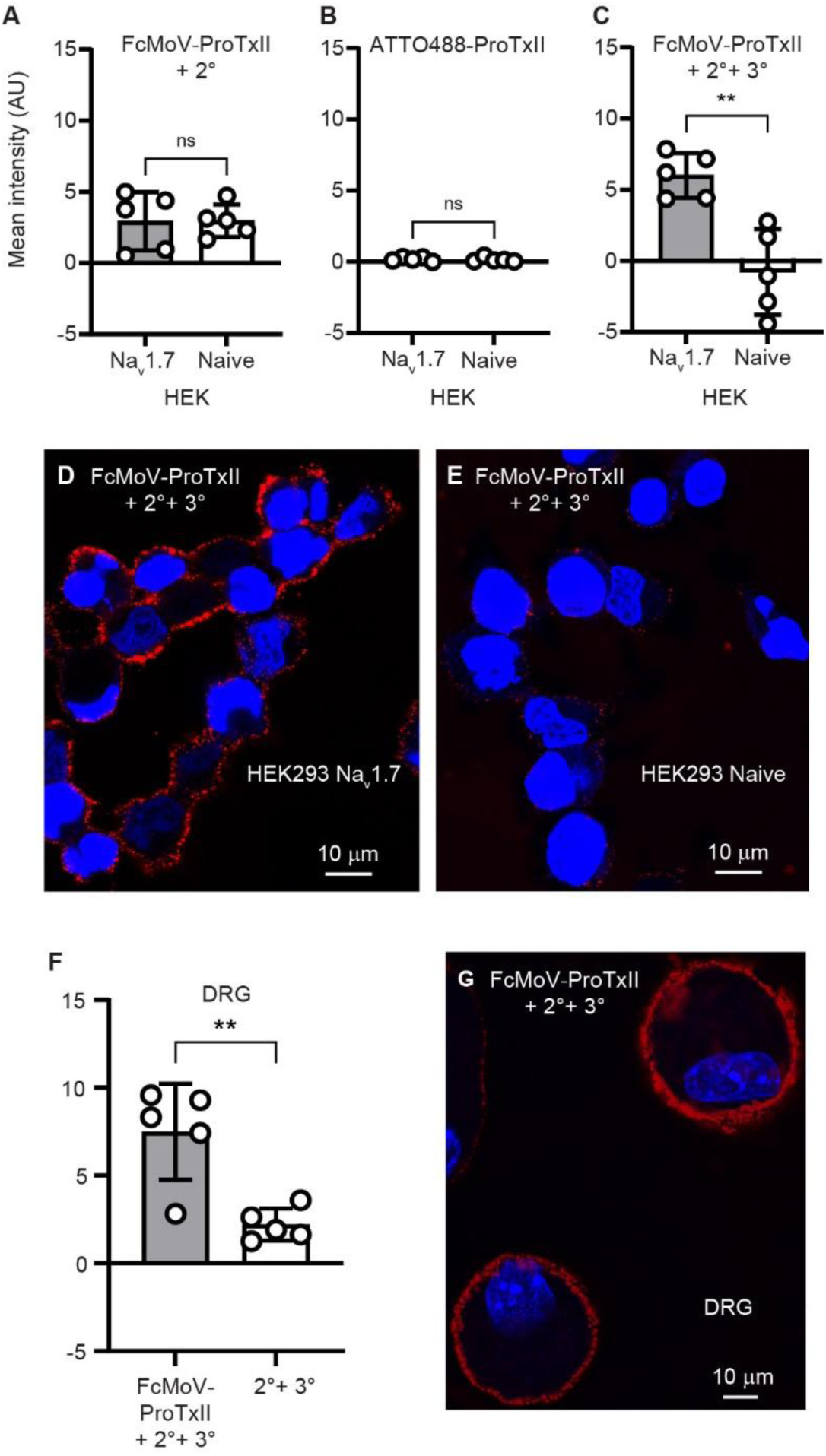
Fluorescent Cell Staining by Monovalent Fc-Toxins A-C. Fluorescence intensity of HEK293 cells either stably expressing Na_v_1.7 or naïve, stained by **A**, FcMoV-ProTxII + secondary antibody AF568 (2°), or **B**, ATTO488-ProTxII, or **C**, FcMoV-ProTxII + secondary and tertiary antibody AF568 (2° + 3°). **D.** Confocal image of HEK293 Na_v_1.7 stable cells labelled with FcMoV-ProTxII + secondary and tertiary antibody AF568 (red). **E.** Same as **D**, but for naïve HEK293 cells. **F.** Fluorescence intensity of DRG neurons stained by FcMoV-ProTxII + secondary and tertiary antibody AF568 versus secondary and tertiary antibody AF568 alone. **G.** Confocal image of DRG neurons labelled with FcMoV-ProTxII + secondary and tertiary antibody AF568 (red). DAPI stained nuclei coloured blue.

## DISCUSSION AND CONCLUSIONS

Toxins rich in disulphide bridges are potent and selective modulators of ion channels, but as we show here, they are typically poorly expressed recombinantly from mammalian cells. We selected a panel of 13 snake, spider and scorpion toxins to assess yields and function when fused to Fc domains. Whilst FcBiV toxin fusions generally improved expression, protein production remained inconsistent. In contrast, FcMoV toxin fusions exhibited robust expression, with 11 of 13 toxins giving final purified yields of greater than 15 μg from 1 ml of media, meaning mg quantities can be obtained from cultures of only 20-70 ml of Expi293F cells. The FcMoV format boosted yields across the panel while simultaneously enabling production of bioactive toxins that would have otherwise eluded functional characterisation due to poor expression, such as JzTxV and ProTxII. We hypothesise that improved FcMoV toxin yields result from the combination of excellent Fc domain expression and improved stability engendered by reducing toxin copy load from two to one. This is highlighted by the observation that FcBiV-HwTxIV but not FcMoV-HwTxIV toxins precipitated during storage at 0°C within weeks of purification. Despite this, the FcMoV format was not sufficient to stabilise expression of highly hydrophobic ant toxins that do not possess disulphide bridges and so will not be useful for every toxin type that exists in nature.

SEC analysis revealed that the fused toxin, despite only representing a small proportion of the total protein mass, exerted strong and unpredictable effects on the monodispersity and aggregation propensity of the Fc fusion constructs. However, FcMoV toxin monodispersity was consistently retained at 100 μM during storage at 0°C for at least one month, and some samples were shown to tolerate snap freeze-thawing at 100 μM, whilst others could tolerate snap-freezing by including 20% glycerol or by reducing the concentration to 5 μM. Furthermore, bioactivity was also retained after being kept at 0°C for one month. These findings support the suitability of the FcMoV toxins for long-term storage and utilisation.

Functional effects were observed for the majority of FcMoV toxins, with maximal effects closely matching synthetic toxins. However, potency was reduced. We hypothesise that the reduction in sensitivity is due, at least in part, to a proportion of the toxins experiencing disulphide scrambling during folding in the recombinant expression system, hence rendering a proportion of the molecules inactive. The FcMoV snake toxins were least affected, with potency losses within one order of magnitude, whereas spider toxins showed two to four orders of magnitude reductions in potency and scorpion toxins showed four orders of magnitude losses in potency. This level of reduction in sensitivity means that some toxins such as HpTxI, which has a low sensitivity even in synthetic form (IC_50_ ∼0.5 μM) (Zhou et al., 2020) do not exhibit activity at doses up to 10 μM, which were the highest tested in this study. The improved retention of potency for snake toxins could suggest that the rigid three-finger motifs provide stricter positioning of the cysteine residues to guide accurate bridge formation. Alternatively, perhaps the more similar mammalian cell expression environment to that of snakes, which are also vertebrates, better supports folding of snake toxins versus toxins from more distantly related species such as arachnids. However, the folding success rate does not appear to be related to bridge density because snake toxin bridge density is ∼1 bridge/15 residues, in between that of spider toxin bridge density at ∼1 bridge/12 residues, and that of scorpion toxin density at ∼1 bridge/16 residues.

In the FcMoV spider toxin selectivity trials, HwTxIV-m3, JzTxV, Pn3a, and ProTxII all exhibited selective inhibition of Na_v_1.7 over other Na_v_ isoforms tested. Notably, three from four of these toxins (JzTxV, Pn3a, and ProTxII) could only be produced with good yield (>15 μg from 1 ml media) in the FcMoV format, rather than FcBiV format. HwTxIV-m3 was marginally selective for Na_v_1.7 over Na_v_1.1 and Na_v_1.6, and selective against Na_v_1.5 and Na_v_1.8. In contrast, JzTxV and Pn3a were fully selective for Na_v_1.7 in the concentration ranges tested, while ProTxII was at least 30-fold selective for Na_v_1.7 over Na_v_1.1, Na_v_1.6, and Na_v_1.8, and selective against Na_v_1.5. These selectivity trends match the literature well in all cases (Schmalhofer et al., 2008; Xiao et al., 2008; Revell et al., 2013; Deuis et al., 2017; Moyer et al., 2018), which suggests, importantly, that despite reduced potency, other pharmacological properties of the toxins, such as native target selectivity, remain unaffected by FcMoV fusion recombinant expression. Thus, the utility of these easy-to-produce fusions as selective pharmacological tools for probing physiological systems remains intact. Furthermore, these agents can be used to fluorescently label channels in cell membranes with a greater sensitivity than synthetic toxins due to polyclonal amplification of the signal.

Overall, this study provides insights into straightforward efficient production of pure, monodisperse antibody-toxin fusions for downstream pharmacological work. The high proportion of expression success with the FcMoV platform could be applicable to many disulphide bridge-rich peptides, and potentially other types beyond those investigated here. Even with suboptimal potency, selectivity remained intact for pharmacological discrimination between targets. In-house toxin production grants researchers control over downstream experiments and establishes the first step in the protein engineering pipeline: the ability to produce protein. The FcMoV platform sets the stage for the future expression of peptide libraries to optimise function, potency, folding, stability, bioactivity, selectivity and staining intensity. In doing so it lowers the barriers to the development of pharmacological tools and therapeutic leads against ion channels, such as the established analgesic target Na_v_1.7.

## ACKNOWLEDGEMENTS

Federico Olivero for DRG culture preparation. P.S.M: BBSRC BB/M024709/1. J.E.G-P: Herchel Smith Fund, Cambridge. J.I.B: MRC MR/W006650/1. A.R, E.St.J.S: joint and equal investment from UKRI and Versus Arthritis (MR/W002426/1) as part of the ADVANTAGE visceral pain consortium through the Advanced Pain Discovery Platform (APDP) initiative. M.D, E.St.J.S: Wellcome Trust Discovery Award (225856/Z/22/Z). S.C.S, A.P.J: British Heart Foundation (PG/19/59/34582 and PG/24/12121). A.S.H, N.S, A.M.R, E.S: Metrion Biosciences Ltd. L.L: Wellcome Trust, Grant Number: 218514/Z/19/Z, Merck Sharp and Dohme Corp. and Janssen Pharmaceutica NV. Y.D: UKRI MRC MR/S007180/1 and MR/Z504099/1.

## AUTHOR CONTRIBUTIONS

S.C.S, Y.D, E.St.J.S, A.P.J, E.S, P.S.M: conceived the research. J.E.G, J.I.B, V.M: protein engineering, expression, purification. A.S.H: Qube electrophysiology. J.I.B: HEK293 immunocytochemistry. S.C.S: Patchliner electrophysiology. N.S, A.M.R: DRG electrophysiology. J.I.B, A.R, M.D: DRG neuron immunocytochemistry. L.L: Ca^2+^ flux assay. J.E.G, P.S.M: wrote the manuscript with input from all other authors.

## AVAILABILITY OF DATA

Data available on request from the authors. FcMoV-toxin DNA plasmids are available from Addgene as follows: pHLsec Fc(Mov) (knob)-AahII - 240560; pHLsec Fc(Mov) (knob)-αBgTx - 240561; pHLsec Fc(Mov) (knob)-αCBTx) - 240562; pHLsec Fc(Mov) (knob)-CITx - 240563; pHLsec Fc(Mov) (knob)-HpTxI - 240564; pHLsec Fc(Mov) (knob)-HwTxIV - 240565; pHLsec Fc(Mov) (knob)-m3-HwTxIV - 240566; pHLsec Fc(Mov) (knob)-JzTxV - 240567; pHLsec Fc(Mov) (knob)-OD1 - 240568; pHLsec Fc(Mov) (knob)-pn3a - 240569; pHLsec Fc(Mov) (knob)-ProTxII - 240570; pHLsec Fc(Mov) (knob)-TsV - 240571; pHLsec Fc(Mov) (knob)-TsVII - 240572; pHLsec Fc(Mov) (knob)-No Insert - 240573; pHLsec Fc Y407T Hole - 240574.

## References

Ahmadi, S., Knerr, J.M., Argemi, L., Bordon, K.C.F., Pucca, M.B., Cerni, F.A., et al. (2020). Scorpion venom: Detriments and benefits. Biomedicines 8:.

Alexandrou, A.J., Brown, A.R., Chapman, M.L., Estacion, M., Turner, J., Mis, M.A., et al. (2016). Subtype-selective small molecule inhibitors reveal a fundamental role for Nav1.7 in nociceptor electrogenesis, axonal conduction and presynaptic release. PLoS One 11:.

Aricescu, A.R., Lu, W., and Jones, E.Y. (2006). A time- and cost-efficient system for high-level protein production in mammalian cells. Acta Crystallogr D Biol Crystallogr 62: 1243–1250.

Beeson, D., Amar, M., Bermudez, I., Vincent, A., and Newsom-Davis, J. (1996). Stable functional expression of the adult subtype of human muscle acetylcholine receptor following transfection of the human rhabdomyosarcoma cell line TE671 with cDNA encoding the ε subunit. Neurosci Lett 207: 57–60.

Bencherif, M., and Lukas, R.J. (1991). Differential Regulation of Nicotinic Acetylcholine Receptor Expression by Human TE671/RD Cells following Second Messenger Modulation and Sodium Butyrate Treatments’. Mol Cell Neurosci 2: 52.

Biswas, K., Nixey, T.E., Murray, J.K., Falsey, J.R., Yin, L., Liu, H., et al. (2017). Engineering Antibody Reactivity for Efficient Derivatization to Generate NaV1.7 Inhibitory GpTx-1 Peptide-Antibody Conjugates. ACS Chem Biol 12: 2427–2435.

Bosmans, F., and Swartz, K.J. (2010). Targeting voltage sensors in sodium channels with spider toxins. Trends Pharmacol Sci 31: 175–182.

Bourne, Y., Talley, T.T., Hansen, S.B., Taylor, P., and Marchot, P. (2005). Crystal structure of a Cbtx-AChBP complex reveals essential interactions between snake α-neurotoxins and nicotinic receptors. EMBO Journal 24: 1512–1522.

Cardoso, F.C., Dekan, Z., Rosengren, K.J., Erickson, A., Vetter, I., Deuis, J.R., et al. (2015). Identification and characterization of ProTx-III [μ-TRTX-Tp1a], a new voltage-gated sodium channel inhibitor from venom of the tarantula Thrixopelma pruriens. Mol Pharmacol 88: 291–303.

Cardoso, F.C., and Lewis, R.J. (2018). Sodium channels and pain: from toxins to therapies. Br J Pharmacol 175: 2138–2157.

Cardoso, F.C., Servent, D., and Lima, M.E. de (2022). Editorial: Venom Peptides: A Rich Combinatorial Library for Drug Development. Front Mol Biosci 9:.

Clairfeuille, T., Cloake, A., Infield, D.T., Llongueras, J.P., Arthur, C.P., Li, Z.R., et al. (2019). Structural basis of a-scorpion toxin action on Nav channels. Science (1979) 363:.

Deuis, J.R., Dekan, Z., Wingerd, J.S., Smith, J.J., Munasinghe, N.R., Bhola, R.F., et al. (2017). Pharmacological characterisation of the highly Na v 1.7 selective spider venom peptide Pn3a. Sci Rep 7:.

Eagles, D.A., Chun, |, Chow, Y., Glenn, |, and King, F. (2020). Fifteen years of Na V 1.7 channels as an analgesic target: Why has excellent in vitro pharmacology not translated into in vivo analgesic efficacy? Br J Pharmacol 179: 3592–3611.

Edwards, W., Fung-Leung, W.P., Huang, C., Chi, E., Wu, N., Liu, Y., et al. (2014). Targeting the ion channel Kv1.3 with scorpion venom peptides engineered for potency, selectivity, and half-life. Journal of Biological Chemistry 289: 22704–22714.

Fang, X.T., Sehlin, D., Lannfelt, L., Syvänen, S., and Hultqvist, G. (2017). Efficient and inexpensive transient expression of multispecific multivalent antibodies in Expi293 cells. Biol Proced Online 19:.

Herzig, V., Cristofori-Armstrong, B., Israel, M.R., Nixon, S.A., Vetter, I., and King, G.F. (2020). Animal toxins — Nature’s evolutionary-refined toolkit for basic research and drug discovery. Biochem Pharmacol 181:.

Kalia, J., Milescu, M., Salvatierra, J., Wagner, J., Klint, J.K., King, G.F., et al. (2015). From foe to friend: Using animal toxins to investigate ion channel function. J Mol Biol 427: 158–175.

Kanellopoulos, A.H., Koenig, J., Huang, H., Pyrski, M., Millet, Q., Lolignier, S., et al. (2018). Mapping protein interactions of sodium channel Na V 1.7 using epitope-tagged gene-targeted mice. EMBO J 37: 427–445.

Kyte, J., and Doolittle, R.F. (1982). A simple method for displaying the hydropathic character of a protein. J Mol Biol 157: 105–132.

Lo, K.M., Sudo, Y., Chen, J., Li, Y., Lan, Y., Kong, S.M., et al. (1998). High level expression and secretion of Fc-X fusion proteins in mammalian cells. Protein Eng 11: 495–500.

Maertens, C., Cuypers, E., Amininasab, M., Jalali, A., Vatanpour, H., and Tytgat, J. (2006). Potent modulation of the voltage-gated sodium channel Nav1.7 by OD1, a toxin from the scorpion Odonthobuthus doriae. Mol Pharmacol 70: 405–414.

Minassian, N.A., Gibbs, A., Shih, A.Y., Liu, Y., Neff, R.A., Sutton, S.W., et al. (2013). Analysis of the structural and molecular basis of voltage-sensitive sodium channel inhibition by the spider toxin huwentoxin-IV (μ-TRTX-Hh2a). Journal of Biological Chemistry 288: 22707–22720.

Montnach, J., Waard, S. De, Nicolas, S., Burel, S., Osorio, N., Zoukimian, C., et al. (2021). Fluorescent- and tagged-protoxin II peptides: potent markers of the Nav1.7 channel pain target. Br J Pharmacol 178: 2632–2650.

Moyer, B.D., Murray, J.K., Ligutti, J., Andrews, K., Favreau, P., Jordan, J.B., et al. (2018). Pharmacological characterization of potent and selective NaV1.7 inhibitors engineered from Chilobrachys jingzhao tarantula venom peptide JzTx-V. PLoS One 13:.

Nagai, T., Ibata, K., Park, E.S., Kubota, M., Mikoshiba, K., and Miyawaki, A. (2002). A variant of yellow fluorescent protein with fast and efficient maturation for cell-biological applications. Nat Biotechnol 20: 87–70.

Neff, R.A., Flinspach, M., Gibbs, A., Shih, A.Y., Minassian, N.A., Liu, Y., et al. (2020). Comprehensive engineering of the tarantula venom peptide huwentoxin-IV to inhibit the human voltage-gated sodium channel hNa v 1.7. Journal of Biological Chemistry 295: 1315–1327.

Nguyen, P.T., Nguyen, H.M., Wagner, K.M., Stewart, R.G., Singh, V., Thapa, P., et al. (2022). Computational design of peptides to target Na V 1.7 channel with high potency and selectivity for the treatment of pain. Elife 1: 1–37.

Oganesyan, V., Gao, C., Shirinian, L., Wu, H., and Dall’Acqua, W.F. (2008). Structural characterization of a human Fc fragment engineered for lack of effector functions. Acta Crystallogr D Biol Crystallogr 64: 700–704.

Pattison, L.A., Rickman, R.H., Hilton, H., Dannawi, M., Wijesinghe, S.N., Ladds, G., et al. (2024). Activation of the proton-sensing GPCR, GPR65 on fibroblast-like synoviocytes contributes to inflammatory joint pain. Proceedings of the National Academy of Sciences 121: e2410653121.

Rahman, M.M., Teng, J., Worrell, B.T., Noviello, C.M., Lee, M., Karlin, A., et al. (2020). Structure of the Native Muscle-type Nicotinic Receptor and Inhibition by Snake Venom Toxins. Neuron 106: 952–962.e5.

Rahnama, S., Deuis, J.R., Cardoso, F.C., Ramanujam, V., Lewis, R.J., Rash, L.D., et al. (2017). The structure, dynamics and selectivity profile of a NaV 1.7 potency-optimised huwentoxin-IV variant. PLoS One 12:.

Revell, J.D., Lund, P.E., Linley, J.E., Metcalfe, J., Burmeister, N., Sridharan, S., et al. (2013). Potency optimization of Huwentoxin-IV on hNav1.7: A neurotoxin TTX-S sodium-channel antagonist from the venom of the Chinese bird-eating spider Selenocosmia huwena. Peptides (N.Y.) 44: 40–46.

Ridgway, J.B.B., Presta, L.G., and Carter, P. (1996). ‘Knobs-into-holes’ engineering of antibody CH3 domains for heavy chain heterodimerization. Protein Eng 9: 617–621.

Robinson, S.D., Deuis, J.R., Touchard, A., Keramidas, A., Mueller, A., Schroeder, C.I., et al. (2023). Ant venoms contain vertebrate-selective pain-causing sodium channel toxins. Nat Commun 14:.

Robinson, S.D., and Vetter, I. (2020). Pharmacology and therapeutic potential of venom peptides. Biochem Pharmacol 181:.

Rush, A.M., Cummins, T.R., and Waxman, S.G. (2007). Multiple sodium channels and their roles in electrogenesis within dorsal root ganglion neurons. Journal of Physiology 579: 1–14.

Salvage, S.C., Rahman, T., Eagles, D.A., Rees, J.S., King, G.F., Huang, C.L.H., et al. (2023). The β3-subunit modulates the effect of venom peptides ProTx-II and OD1 on NaV1.7 gating. J Cell Physiol 238: 1354–1367.

Schindelin, J., Arganda-Carreras, I., Frise, E., Kaynig, V., Longair, M., Pietzsch, T., et al. (2012). Fiji: An open-source platform for biological-image analysis. Nat Methods 9: 676– 682.

Schmalhofer, W.A., Calhoun, J., Burrows, R., Bailey, T., Kohler, M.G., Weinglass, A.B., et al. (2008). ProTx-II, a selective inhibitor of NaV1.7 sodium channels, blocks action potential propagation in nociceptors. Mol Pharmacol 74: 1476–1484.

Sermadiras, I., Revell, J., Linley, J.E., Sandercock, A., and Ravn, P. (2013). Recombinant expression and in Vitro characterisation of active huwentoxin-IV. PLoS One 8:.

Wang, R.E., Wang, Y., Zhang, Y., Gabrelow, C., Zhang, Y., Chi, V., et al. (2016). Rational design of a Kv1.3 channel-blocking antibody as a selective immunosuppressant. Proc Natl Acad Sci U S A 113: 11501–11506.

Wulff, H., Christophersen, P., Colussi, P., Chandy, K.G., and Yarov-Yarovoy, V. (2019). Antibodies and venom peptides: new modalities for ion channels. Nat Rev Drug Discov 18: 339–357.

Xiao, Y., Bingham, J.P., Zhu, W., Moczydlowski, E., Liang, S., and Cummins, T.R. (2008). Tarantula huwentoxin-IV inhibits neuronal sodium channels by binding to receptor site 4 and trapping the domain II voltage sensor in the closed configuration. Journal of Biological Chemistry 283: 27300–27313.

Xu, H., Li, T., Rohou, A., Arthur, C.P., Tzakoniati, F., Wong, E., et al. (2019). Structural Basis of Nav1.7 Inhibition by a Gating-Modifier Spider Toxin. Cell 176: 702–715.

Zeisel, A., Hochgerner, H., Lönnerberg, P., Johnsson, A., Memic, F., Zwan, J. van der, et al. (2018). Molecular Architecture of the Mouse Nervous System. Cell 174: 999–1014.e22.

Zhou, X., Ma, T., Yang, L., Peng, S., Li, L., Wang, Z., et al. (2020). Spider venom-derived peptide induces hyperalgesia in Nav1.7 knockout mice by activating Nav1.9 channels. Nat Commun 11:.

