## Supplemental Figs 1-5 for "A robust platform for recombinant production of animal venom toxin modulators of ion channels"

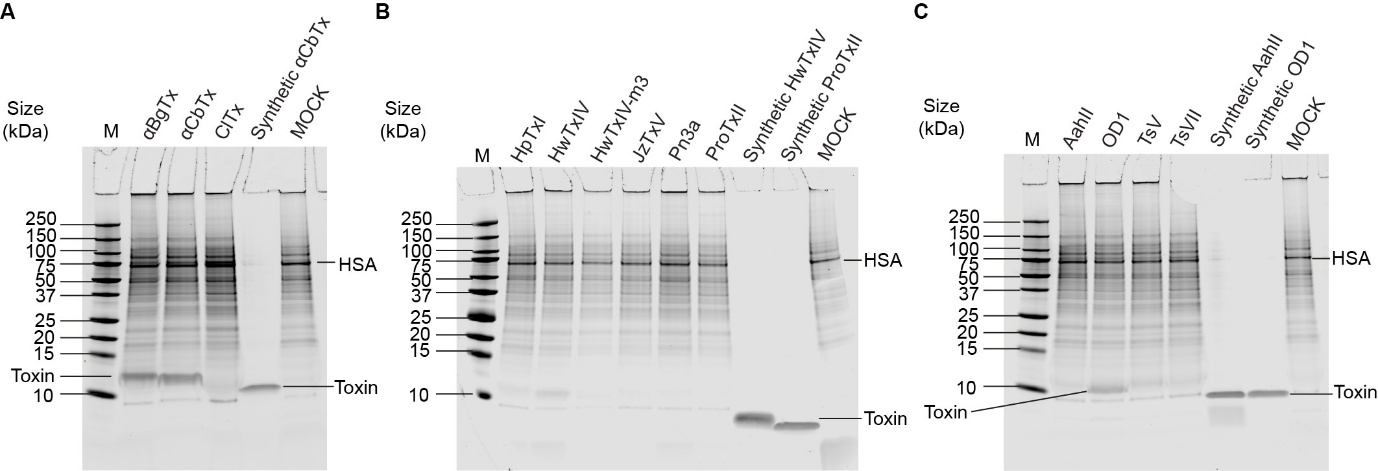

**FIGURE S1 – Expression Trial of Unfused Peptide Toxins**

13 toxins were transfected into Expi293F cells, and the resultant media supernatant (cells removed; 40 μl loads) was run on Coomassie stained reducing SDS-PAGE alongside synthetic toxin positive controls (1 μg loads) and mock transfection negative controls. **A**. Snake toxin gel showing expression bands for αBgTx and αCbTx (2 of 3 toxins). **B.** Spider toxin gel showing no successful expression (0 of 6 toxins). **C.** Scorpion toxin gel showing an expression band for OD1 (1 of 4 toxins). M = size marker; HSA = human serum albumin; MOCK = mock transfection with no toxin DNA.

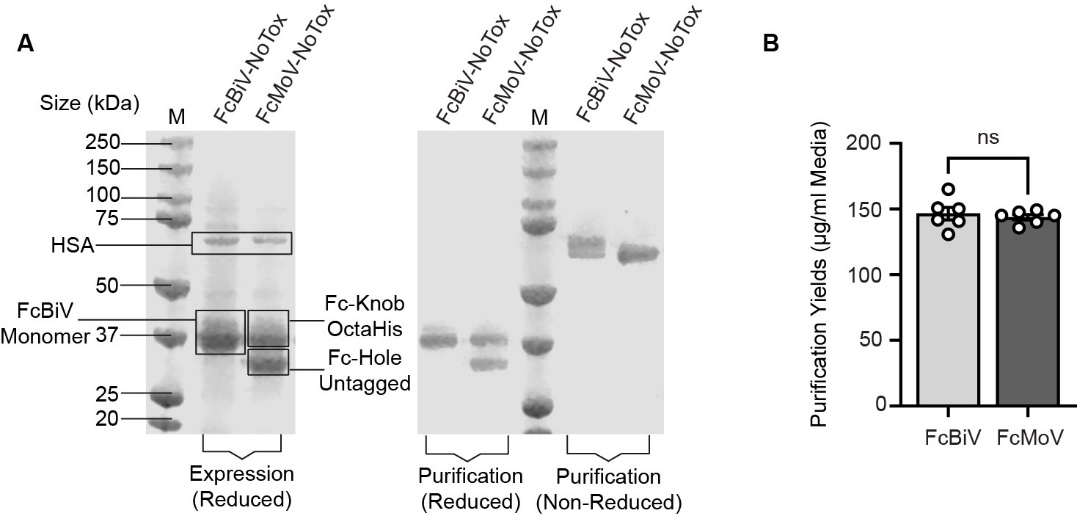

**FIGURE S2 – Expression Trial of Unfused FcBiV & FcMoV Constructs**

To investigate which of the Fc formats, FcBiV or FcMoV, yielded more protein without any fused toxin, unfused FcBiV homodimers and FcMoV heterodimers were expressed from Expi293F cells, purified using Ni-NTA agarose beads, and yields were quantified. **A.** Coomassie stained SDS-PAGE gels of expression media supernatant (left, reducing) and purified protein (right, reducing versus non-reducing), revealing strong expression bands for both constructs, as well as high purity and intact dimerisation states. **B.** Bar chart showing purification yields of unfused FcBiV versus FcMoV, µg per ml of expression media (n = 6; Unpaired t-test, two-tailed; ns > 0.05). M = size marker; HSA = human serum albumin.

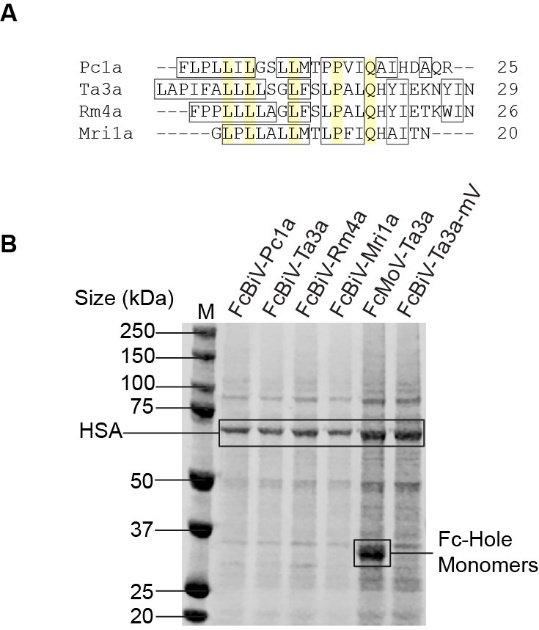

**FIGURE S3 – Expression Trial of Ant Fc-Toxin Fusions**

**A**. Protein sequence alignment of ant peptide toxins: Pc1a, Ta3a, Rm4a, and Mri1a. Conserved residues and highlighted in yellow; hydrophobic residues are shown inside black boxes. **B**. Coomassie stained reducing SDS-PAGE gel of Expi293F media supernatant (cells removed; 10 μl loads) from recombinant expression trial. None of the four FcBiV ant toxins were successfully expressed from Expi293F cells. One of the toxins, Ta3a, was tested in three different formats: FcBiV, FcMoV, and FcBiV with C-terminal fusion to a monoVenus (mV) fluorophore; however, none of these formats successfully rescued recombinant expression. M = size marker; HSA = human serum albumin.

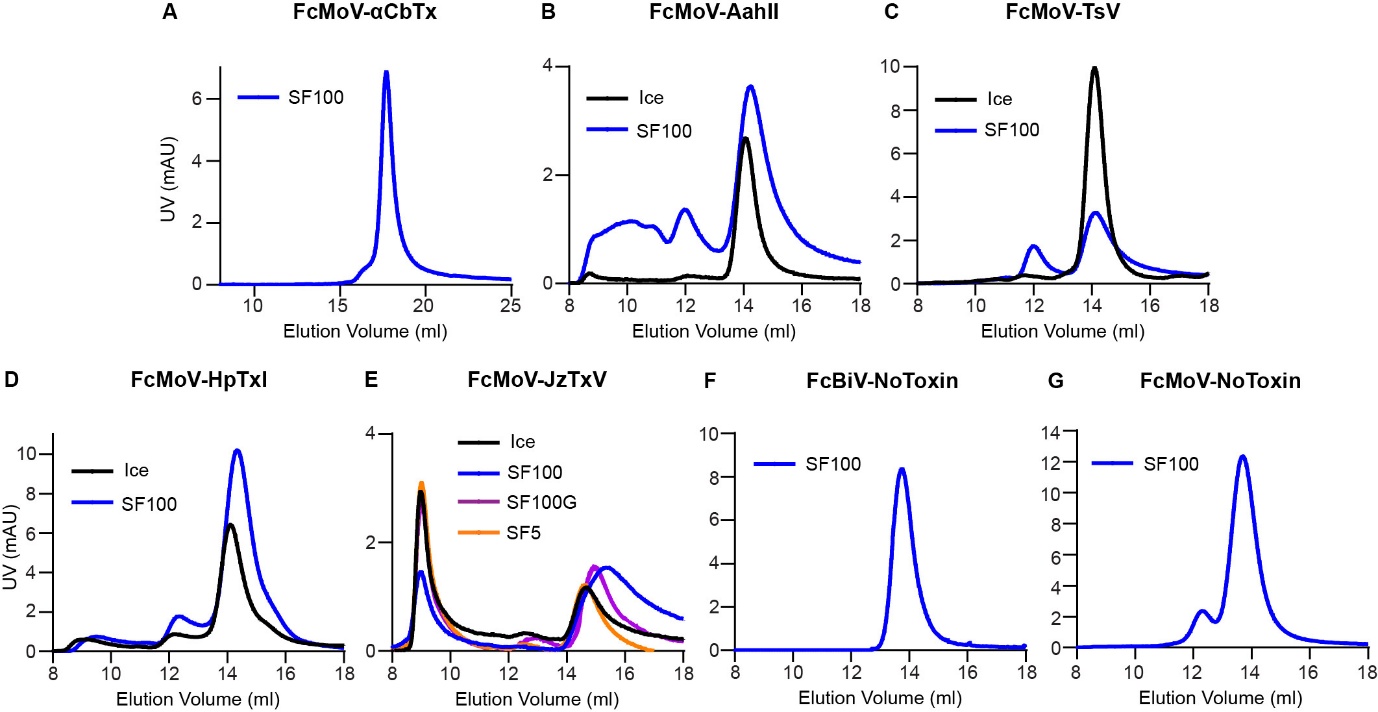

**FIGURE S4 – Additional Fc-Toxin, Unfused FcBiV & Unfused FcMoV SEC Traces**

All purified FcMoV toxins, and the highest-yielding purified FcBiV toxin (HwTxIV), were run on size-exclusion chromatography (SEC) using 70 µg loads to explore protein monodispersity and optimal storage conditions (n = 1 run per toxin condition; n = 28 total SEC runs). Ultraviolet (UV) absorbance peaks at 14-15 ml elution volume represented the ~70 kDa monomeric Fc toxin species size, indicative of high protein quality. A higher order aggregation (agg) peak at 8-10 ml elution volume, or secondary (2⁰) shoulder peak at 11-13 ml elution volume were sometimes also visible. SEC trace colours: black (Ice) = stored on ice; blue (SF100) = snap-freeze thawed at 100 μM; purple (SF100G) = snap freeze-thawed at 100 µM with 20% glycerol; orange (SF5) = snap freeze-thawed at 5 µM. **A.** FcMoV-αCbTx trace. **B**. FcMoV-AahII traces. **C**. FcMoV-TsV traces. **D.** FcMoV-HpTxI traces. **E**. FcMoV-JzTxV traces. **F**. Unfused FcBiV homodimer trace. **G**. Unfused FcMoV heterodimer trace.

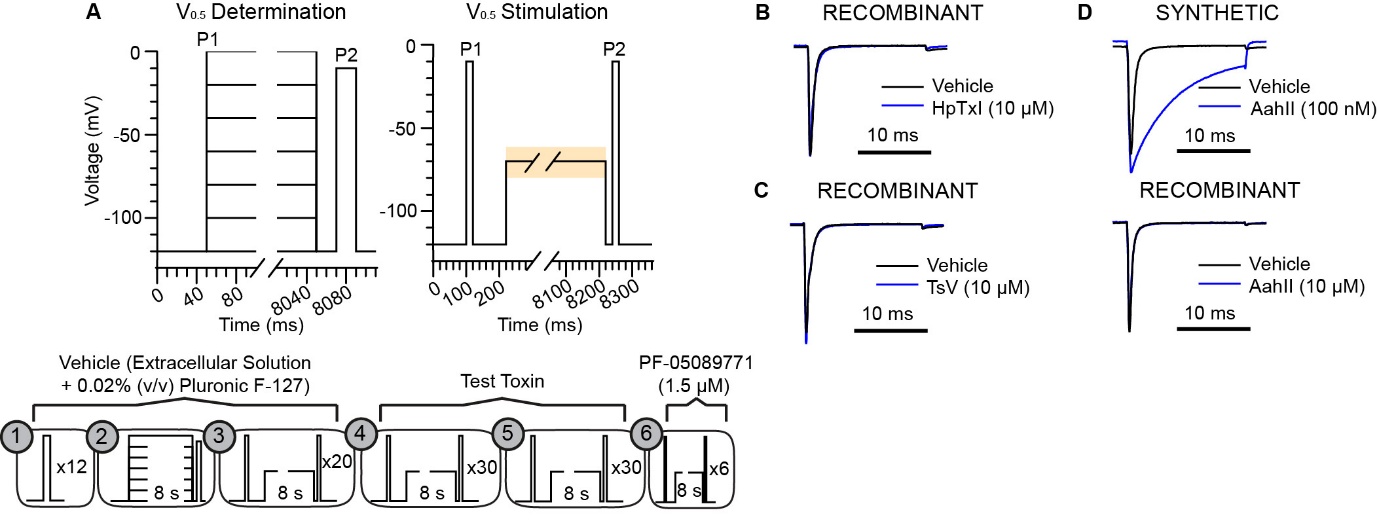

**FIGURE S5 – Additional Monovalent Fc-Toxin Electrophysiology Information**

**A.** Voltage protocol for high-throughput automated electrophysiology using the Qube. Independent cells were subjected to an I-V (current-voltage) protocol to determine the voltage that elicits half-channel (V_0.5_) inactivation using fractional recovery of P2 versus P1 (top, left). This V_0.5_ inactivation value, unique to each cell, was differentially applied in the screening protocol used to assess toxin activity (top, right; the orange box denotes V_0.5_ inactivation implementation). The assay involved six phases (1-6; bottom): 1 = initial current stabilisation; 2 = V_0.5_ determination (3 min); 3 = V_0.5_ stimulation in vehicle to obtain a stable baseline (10 min); 4 & 5 = V_0.5_ stimulation in toxin (2 x 15 min); 6 = V_0.5_ stimulation in positive control PF-05089771 for full current inhibition (3 min). **B.** Whole-cell patch clamp recordings of Na^+^ currents in CHO cells stably expressing Na_V_1.7 in the absence (black trace) and presence (blue trace) of recombinant FcMoV HpTxI. **C.** The same but for FcMoV-TsV. **D.** The same but for synthetic AahII (top panel) and FcMoV-AahII (bottom panel).
