## Supplementary material for "A robust platform for recombinant production of animal venom toxin modulators of ion channels": Table 1

| **Toxin Name (Abbreviation)** | **Animal of Origin** | **Species** | **Main Mechanism of Action** | **Reference** |
| --- | --- | --- | --- | --- |
| α-Bungarotoxin (αBgTx) | Snake | *Bungarus multicinctus* | nAChR Inhibitor | (Rahman et al., 2020) |
| α-Cobratoxin (αCbTx) |  | *Naja kaouthia* | nAChR Inhibitor | (Bourne et al., 2005) |
| Calliotoxin (ClTx) |  | *Calliophis bivirgatus* | Na_v_ Potentiator | (Yang et al., 2016) |
| *Heteropoda venatoria* toxin I (HpTxI) | Spider | *Heteropoda venatoria* | Na_v_ Inhibitor | (Zhou et al., 2020) |
| Huwentoxin-IV (HwTxIV) |  | *Ornithoctonus huwena* | Na_v_ Inhibitor | (Minassian et al., 2013) |
| Huwentoxin-IV-m3 (HwTxIV-m3) |  | *Ornithoctonus huwena* | Na_v_ Inhibitor | (Rahnama et al., 2017) |
| Jingzhaotoxin-V (JzTxV) |  | *Chilobrachys jingzhao* | Na_v_ Inhibitor | (Moyer et al., 2018) |
| µ-Theraphotoxin-Pn3a (Pn3a) |  | *Pamphobeteus nigricolor* | Na_v_ Inhibitor | (Deuis et al., 2017) |
| Protoxin-II (ProTxII) |  | *Thrixopelma pruriens* | Na_v_ Inhibitor | (Schmalhofer et al., 2008) |
| *Androctonus australis* *hector* toxin II (AahII) | Scorpion | *Androctonus australis* | Na_v_ Potentiator | (Alami et al., 2003) |
| *Odonthobuthus doriae* toxin 1 (OD1) |  | *Odonthobuthus doriae* | Na_v_ Potentiator | (Maertens et al., 2006) |
| *Tityus serrulatus* toxin V (TsV) |  | *Tityus serrulatus* | Na_v_ Potentiator | (Pucca et al., 2015a) |
| *Tityus serrulatus* toxin VII (TsVII) |  | *Tityus serrulatus* | Na_v_ Potentiator | (Pucca et al., 2015b) |

**Table 1** – Toxin Information: Animal Origin & General Mechanism of Action
