## Supplementary material for "A robust platform for recombinant production of animal venom toxin modulators of ion channels": Table 2

**Table 2** – Concentration of Spider Toxins for Half-Maximal Inhibition of Five Na_v_ Isoforms

| FcMoV Spider Toxin | IC_50_ (nM) | | | | |
| --- | --- | --- | --- | --- | --- |
|  | Na_v_1.7 | Na_v_1.1 | Na_v_1.5 | Na_v_1.6 | Na_v_1.8 |
| HwTxIV-m3 | 102 ± 27 | 684 ± 188 | >10,000 | 670 ± 133 | >10,000 |
| JzTxV | 938 ± 241 | >10,000 | >10,000 | >10,000 | >10,000 |
| Pn3a | 1532 ± 178 | >10,000 | >10,000 | >10,000 | >10,000 |
| ProTxII | 242 ± 68 | 7693 ± 2052 | >10,000 | >10,000 | 7673 ± 909 |
